# Protocell Arrays for Simultaneous Detection of Diverse Analytes

**DOI:** 10.1101/2021.02.13.431022

**Authors:** Yan Zhang, Taisuke Kojima, Ge-Ah Kim, Monica P. McNerney, Shuichi Takayama, Mark P. Styczynski

## Abstract

Simultaneous detection of multiple analytes from a single sample (multiplexing), particularly when at the point of need, can guide complex decision-making without increasing the required sample volume or cost per test. Despite recent advances, multiplexing still typically faces the critical limitation of measuring only one type of molecule per assay platform – for example, only small molecules or only nucleic acids. In this work, we address this bottleneck with a customizable platform that integrates cell-free expression (CFE) with a polymer-based aqueous two-phase system (ATPS) to produce membrane-less “protocells” containing transcription and translation machinery used for analyte detection. Multiple protocells are arrayed in microwells where each protocell droplet performs distinct reactions to detect chemically diverse targets including small molecules, minerals, and nucleic acid sequences, all from the same sample. We demonstrate that these protocell arrays can measure analytes in a human biofluid matrix, maintain function after lyophilization and rehydration, and produce visually interpretable readouts, illustrating its potential for application as a minimal-equipment, field-deployable, multi-analyte detection tool.

## Introduction

The ability to sense and characterize the concentrations of analytes is key to many engineering and biomedical advances, especially when these measurements can be made at or near the point of need. For example, rapid measurement of analyte concentrations in chemical processes can enable external control of those processes or detect faults or dangerous conditions that must be addressed. While this type of analysis is sometimes straightforward to perform for measurement of a single target, many samples with industrial or biomedical relevance have complex analyte profiles and often require measurement of multiple targets spanning different molecular classes (from ions to small molecules to nucleic acids). Current sensing and diagnostic tools, especially those designed for use at the point of need, typically only detect one type of analyte at a time, which places significant limitations on the number and the type of tests that can be done on each sample^1^. Thus, being able to simultaneously detect multiple analytes across different molecular classes from a single sample at the point-of-need will empower researchers and clinicians with more information without requiring additional assay time, sample volume, and cost per test.

Unlike man-made sensors and diagnostics, cells are equipped with extensive capabilities to simultaneously detect multiple, diverse types of analytes. Recently developed “cell-free” biosensors, where cell lysates are used as rich reaction mixtures, have taken advantage of some of this cellular machinery to sense and respond to target analytes *via* protein or RNA expression^2^. The component reagents can also be lyophilized to enable wide-scale, affordable, and deployable testing^2^. Sensors based on cell-free expression (CFE) have been used for point-of-need testing of environmental contaminants for problems including water quality monitoring^3–5^. CFE-based sensors have also been used to detect clinically relevant targets with minimal equipment and operator requirements^6–8^, making them excellent candidates for widespread use. However, nearly all current CFE sensors can detect only one analyte, even though the sample assayed may contain multiple relevant analytes. To detect all relevant analytes, multiple CFE reactions would need to be run in parallel using multiple aliquots of the sample, introducing handling challenges for onsite testing and collection challenges for samples that are often only available in small volumes. Existing strategies for multiplexed testing – which is the simultaneous detection of different analytes from a single sample – often require a library of orthogonal reporters, cannot measure analytes from diverse molecular classes (i.e., both nucleic acid sequences and small molecules), or cannot be used in minimal-equipment settings^1,9–13^. To effectively and efficiently address impactful problems such as the diagnosis of diseases with complex biomarker profiles at the point of need, it is critical to develop improved platforms for multiplexed analysis.

Arrays of membrane-less protocell sensors formed by polymer-based aqueous two-phase systems (ATPS) and CFE reactions have the potential to address current limitations in point-of-need measurement multiplexing. Mixing two immiscible aqueous polymer solutions can lead to spontaneous liquid-liquid phase separation, which in appropriate ratios yields “droplets” in a “bulk phase”. Adding biological machinery (like that of a CFE reaction) to the droplets yields what is essentially a membrane-less precursor of a cell (a “protocell”)^14,15^, with genetic information and the capability to execute complex functions contained in a small localized volume. Furthermore, since many soluble macromolecules such as the proteins and nucleic acids that enable CFE-based sensing selectively partition from a polyethylene glycol (PEG)-rich phase to the dextran- or Ficoll-rich phase^16^, the core of distinct biosensing machinery can stay compartmentalized in each ATPS-formed protocell. Because these protocells are membrane-less, this approach also resolves challenges in transporting macromolecular analytes from the bulk phase to protocells present in most membrane-based approaches^17^. For multiplexed detection of target analytes present from a single sample, we use simple topographical placement of micro-basins inside a microwell for stable positioning of distinct protocells^18,19^, each containing a different CFE sensor (**Figure 1, S1**). Addition of a sample solution to such a microwell triggers analyte uptake and compartmentalized detection reactions in multiple isolated membrane-less protocells.

**Figure 1:**
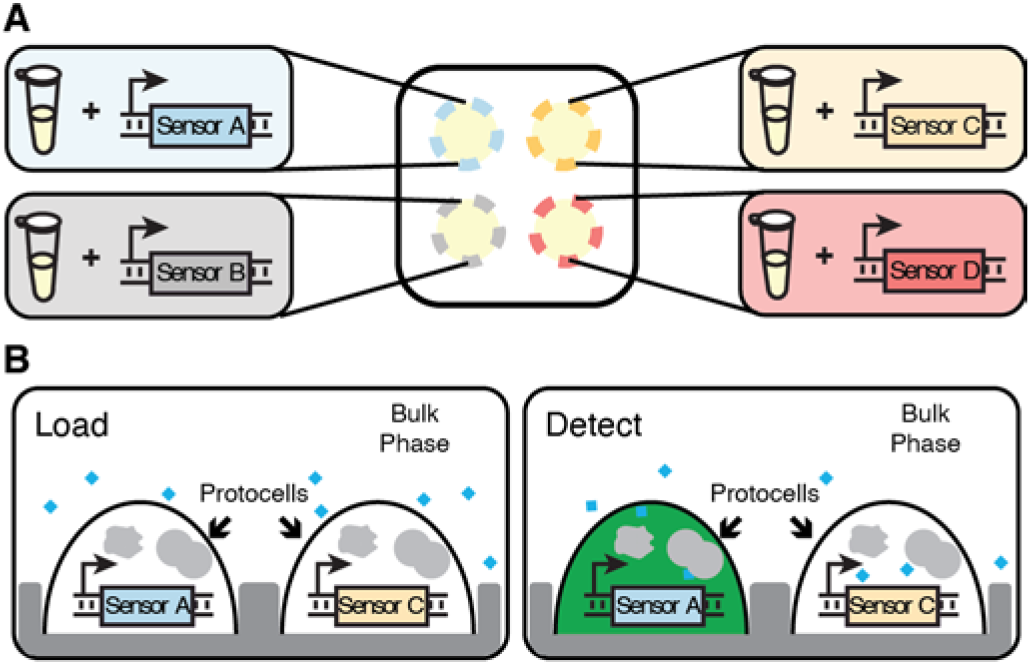
Schematic of the membrane-less protocell array-based, multiplexed, multiple analyte sensing platform. The use of spatially patterned arrays of phase-separated protocell sensors enables each microwell to provide measurements for multiple analytes using a single type of fluorescent or colorimetric readout. **(A)** An overhead view of a microwell with four micro-basins. White area surrounding the micro-basins (yellow-shaded regions) represents the bulk phase formed by a PEG solution containing the analytes to be detected. Each micro-basin has a protocell containing CFE reaction (represented by the microcentrifuge tube) with a different sensor plasmid or genetic circuit. **(B)** Side view of a microwell with micro-basins. Analytes in the bulk phase (blue diamonds) enter protocells to activate their cognate sensor (blue DNA construct), producing detectable reporter signals. Levels of multiple analytes can be simultaneously assessed by tracking which protocell produces reporter signals (green color on the right).

To date, simple protocells formed by PEG and dextran or coacervates have been successfully used to study membrane-less compartmentalization of cell lysates and biomolecules^14,15,20^. Recent studies have also shown that ATPS droplets comprised of dehydrated polymers, including proteins, are compatible with storage at ambient temperature^21,22^, enabling low-cost distribution of ATPS immunoassay-based diagnostics to the point of need without cold chain transportation. We envision that arrays of membrane-less protocell sensors formed by selective compartmentalization of CFE reactions in ATPS can facilitate multiplexed analyte detection beyond just proteins, leading to a new class of protocell array-based diagnostics that reports on diverse types of analytes, has a high degree of sensor customizability, and can be used at the point of need.

Toward this goal, here we first assess CFE transcription and translation capabilities in two protocell-forming polymer ATPS environments (PEG-Ficoll and PEG-dextran). We show that analytes (small molecules and nucleic acids) added to a microwell containing an array of CFE protocells selectively activate their cognate sensors confined to distinct protocells with comparable sensitivity to those in non-protocell CFE reactions. We highlight the utility of this platform for simultaneous, multi-modal biomarker detection by demonstrating detection of clinically relevant targets (e.g. nucleic acids from pathogenic bacteria and micronutrients) from the same sample and in a human serum matrix. Only one reporter (green fluorescent protein, GFP) is needed for multiplexing, which significantly reduces the complexity of test development and reconfiguration. These protocell arrays also meet key criteria for field-deployable sensors and diagnostics: we demonstrate that the GFP reporter can be easily replaced with color-producing enzymatic reporters to enable equipment-free test interpretation, and we demonstrate multiplexed sensing using protocell arrays that have been lyophilized for storage at ambient temperature. Taken together, the presented multiplexing platform both expands the reach of cell-free biosensors and provides the foundation to develop advanced, customizable “diagnostic chips” for simultaneous detection of diverse types of analytes at the point of need.

## Results

### Compartmentalization of Cell-Free Protein Expression in ATPS-formed Protocells

While CFE protocells have previously been described^14,20^, their application to analyte detection has not. To use membrane-less CFE protocells for multiplexed analyte measurement, the sensor plasmids and core machinery for transcription and translation must (1) retain activity in an ATPS environment and (2) stay compartmentalized in the protocell. We first verified that CFE reactions maintain transcription and translation function in the context of an ATPS. CFE reactions that constitutively produce GFP were tested in two ATPS environments: a PEG-dextran system and a PEG-Ficoll system. For both ATPS environments, PEG constitutes the bulk phase, and dextran or Ficoll constitutes the protocell **(Figure 2A)**. Individual protocells were first formed by mixing CFE reactions with Ficoll or dextran polymer and adding to a bulk phase PEG solution. No protein production was observed after 3 hours of incubation at 37 °C (**Figure 2B**), perhaps due to necessary salts and building blocks for protein synthesis (nucleotides and amino acids originally in the protocell) not remaining strongly partitioned in the protocell but instead diffusing into the bulk phase and thus diluting their concentrations. By supplementing CFE reaction buffer in the PEG bulk phase, compartmentalized CFE protein synthesis was enabled (**Figure 2B**).

**Figure 2:**
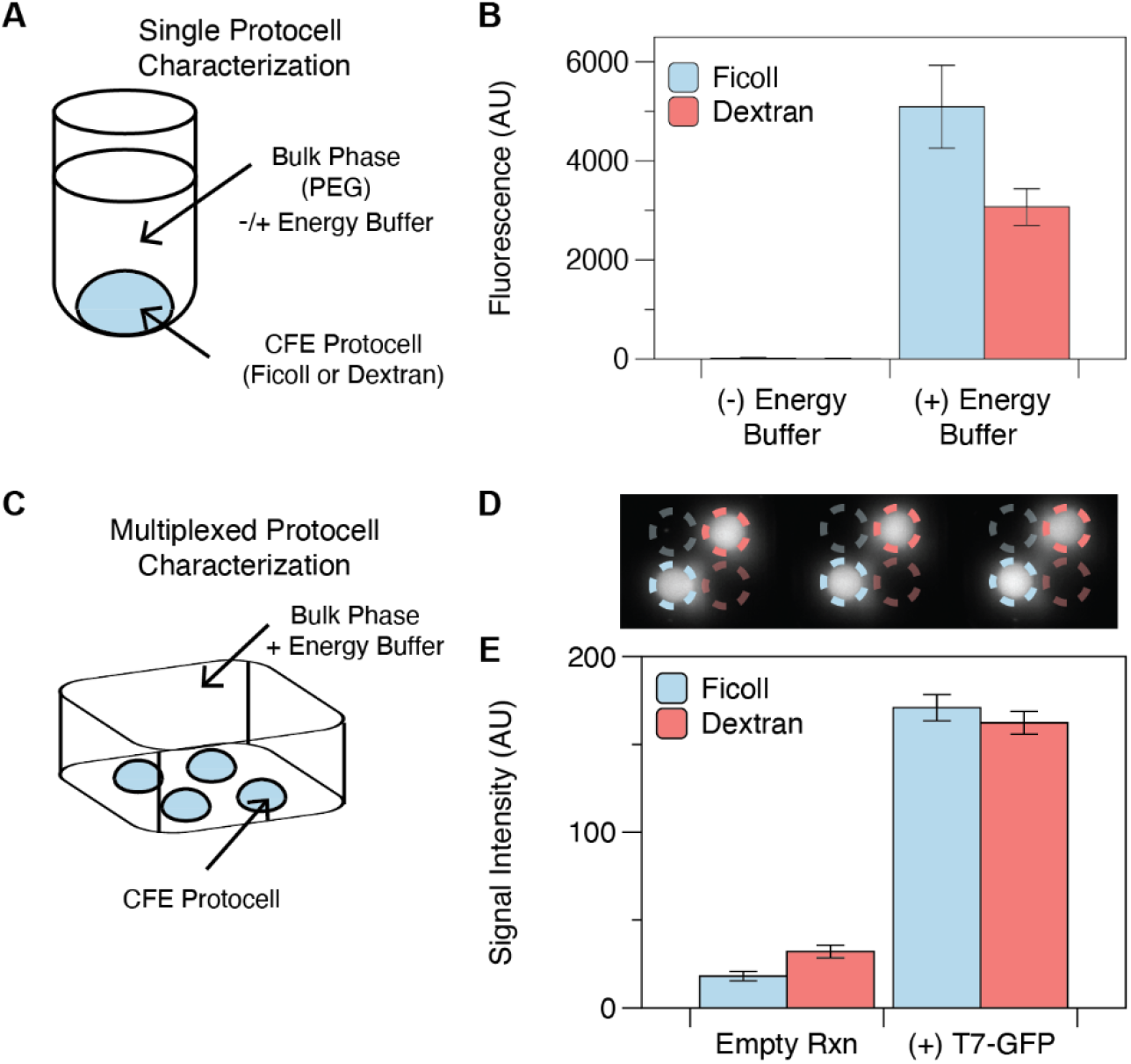
Characterization of protein expression and reaction compartmentalization in membrane-less protocells. **(A)** Schematic of a single protocell reaction. To mimic the final multiplexing condition, a protocell (formed by Ficoll or dextran) expressing GFP was submerged in bulk phase solution. **(B)** Fluorescence (measured with a plate reader) of Ficoll and dextran protocell reactions in single-protocell format. For detectable protein expression, energy buffer must be supplemented in the bulk phase. Error bars represent standard deviations of technical triplicates. **(C)** Schematic of multiplexed protocell array system. Protocells containing different CFE reactions were loaded into micro-basins and submerged in the bulk phase solution. **(D)** Fluorescence image of protocell arrays obtained by a ChemiDoc MP imager. Contents of individual protocells are indicated with colored circles: blue for Ficoll polymer-encapsulated reactions, red for dextran-encapsulated reactions. Bright colors indicate CFE reactions containing GFP plasmid for expression, and faded colors indicate negative controls with no plasmid. **(E)** Quantitative assessment of fluorescence image in (D), with pixel intensity quantified by image processing software (Fiji). Error bars represent standard deviations of technical triplicates. Note, due to different measurement instruments being used for (B) and (E), the arbitrary units on the y-axes are different scales.

We next validated that ATPS-formed protocells compartmentalize multiple CFE reactions, such that reporter plasmids and proteins remain in their own phase-separated droplets in a membrane-less protocell array. We performed CFE reactions in a custom-designed 96-well plate where each microwell contains four micro-basins for arraying protocells (**Figure S1A**) amidst a bulk phase supplemented with energy buffer. CFE reactions with or without a plasmid coding for GFP were combined with dextran or Ficoll, loaded into the micro-basins (**Figure 2C**), and incubated. To measure fluorescence in this custom microwell plate, we used a fluorescence imager (ChemiDoc MP), yielding the desired compartmentalized reaction data from our protocell array. All protocells remained localized in their designated micro-basins with minimal signal crosstalk between empty and GFP-expressing reactions (**Figures 2D, E**), demonstrating that cell-free lysate, plasmids, and reporter proteins remain compartmentalized. This result suggests that multiple protocells expressing different biosensors could be arrayed in the same microwell, enabling a multiplexing platform. Although both dextran- and Ficoll-based protocells showed similar terminal protein production and compartmentalization efficiency, we used Ficoll for all subsequent experiments based on its slightly faster protein production (**Figure S2**).

### Multiplexed Detection of Model Small Molecule and Nucleic Acid Systems

To demonstrate that these membrane-less CFE protocell arrays can be used for multiplexed analyte detection, we next incorporated sensors that respond to multiple model small molecules. Plasmids encoding each sensor system were embedded in different protocells, with analytes added to the bulk phase that should diffuse into the protocells to activate their cognate sensor proteins. To evaluate small molecule detection, we tested the isopropyl β-D-1-thiogalactopyranoside (IPTG)-inducible LacI-P_T7LacO_ system and the arabinose-inducible AraC-P_BAD_ system (**Figure 3A**)^23^. In each system, the small molecule inducer modulates the activity of its corresponding transcriptional regulator (IPTG derepresses LacI and arabinose activates AraC), allowing transcription of GFP reporter from its respective promoter (P_T7LacO_ or P_BAD_). The arabinose CFE sensor uses a single plasmid encoding constitutive AraC expression and P_BAD_ - regulated GFP expression from the same divergent promoter, and the IPTG sensor uses one plasmid to express GFP from a P_T7LacO_ promoter and another to express LacI constitutively. When IPTG and arabinose are individually added to the bulk phase, only the appropriate sensor reactions are activated, with minimal signal crosstalk (**Figures 3B, C**). This demonstrates that protocell arrays can successfully multiplex measurement of small molecule signals.

**Figure 3:**
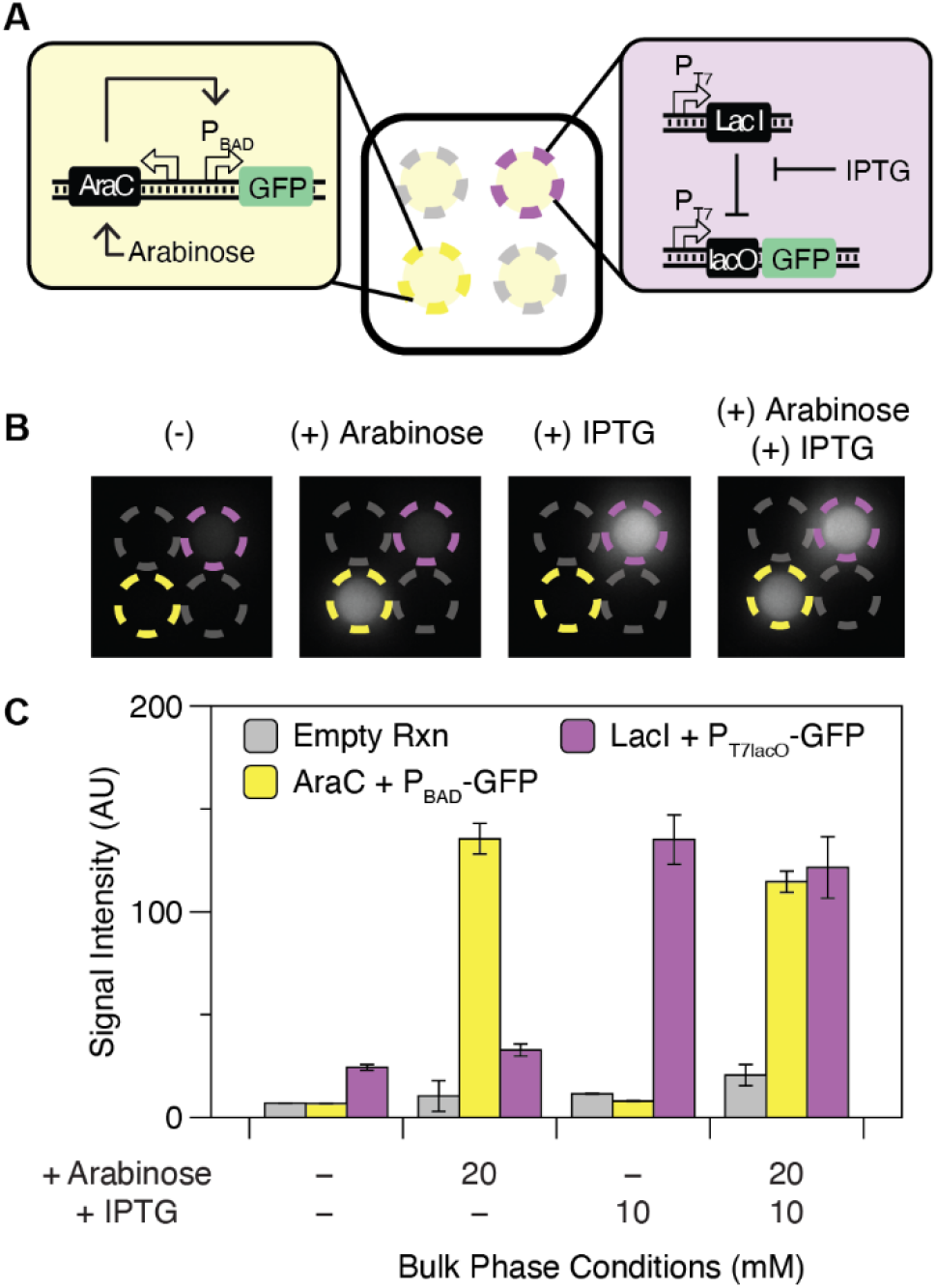
Multiplexed detection of model small molecules in membrane-less protocell arrays. **(A)** Schematic of protocell array reaction setup for multiplexed analysis of small molecules. Yellow and purple circles indicate micro-basins containing arabinose and IPTG sensors, respectively, while gray circles are micro-basins with no-plasmid control protocells. The circuit diagrams for the arabinose and IPTG sensors are shown in the insets. **(B)** Representative fluorescence images of results of multiplexed measurement of model small molecules. Small molecules added to the bulk phase for each experiment are indicated above each image. Circle colors are as in (A). Each protocell sensor is only activated when its cognate small molecule is present in the bulk phase. **(C)** Quantification of fluorescence images in (B) and their replicates. Error bars represent standard deviations of technical triplicates.

We next demonstrated that protocell arrays can also multiplex the detection of RNA targets using previously-reported model toehold switches to control the output of CFE reactions^24^. A toehold switch is a *de novo* designed RNA regulator that forms an inhibitory hairpin to prevent translation of a downstream protein^25^. Addition of a “trigger” RNA with sequence complementarity to part of the switch unfolds the hairpin, turning on reporter expression (**Figure 4A**). We constructed plasmids in which two previously characterized orthogonal toehold switches (B and H)^24^ are constitutively expressed from a T7 promoter, and added these sensor plasmids to the protocell array (**Figure 4B**). Different RNA triggers were added to the bulk phase; GFP expression for a given protocell sensor was only observed when the cognate RNA trigger for the switch in that protocell was present in the bulk phase, demonstrating that protocell arrays can successfully multiplex detection of RNA targets (**Figures 4C, D**). We also note that sensing reactions in protocell arrays had slightly improved limits of detection and fold-activation at low trigger concentrations compared to single-phase CFE reactions (**Figure S3**), likely due to selective RNA partitioning into the protocell-forming phase causing increased local concentrations^15^.

**Figure 4:**
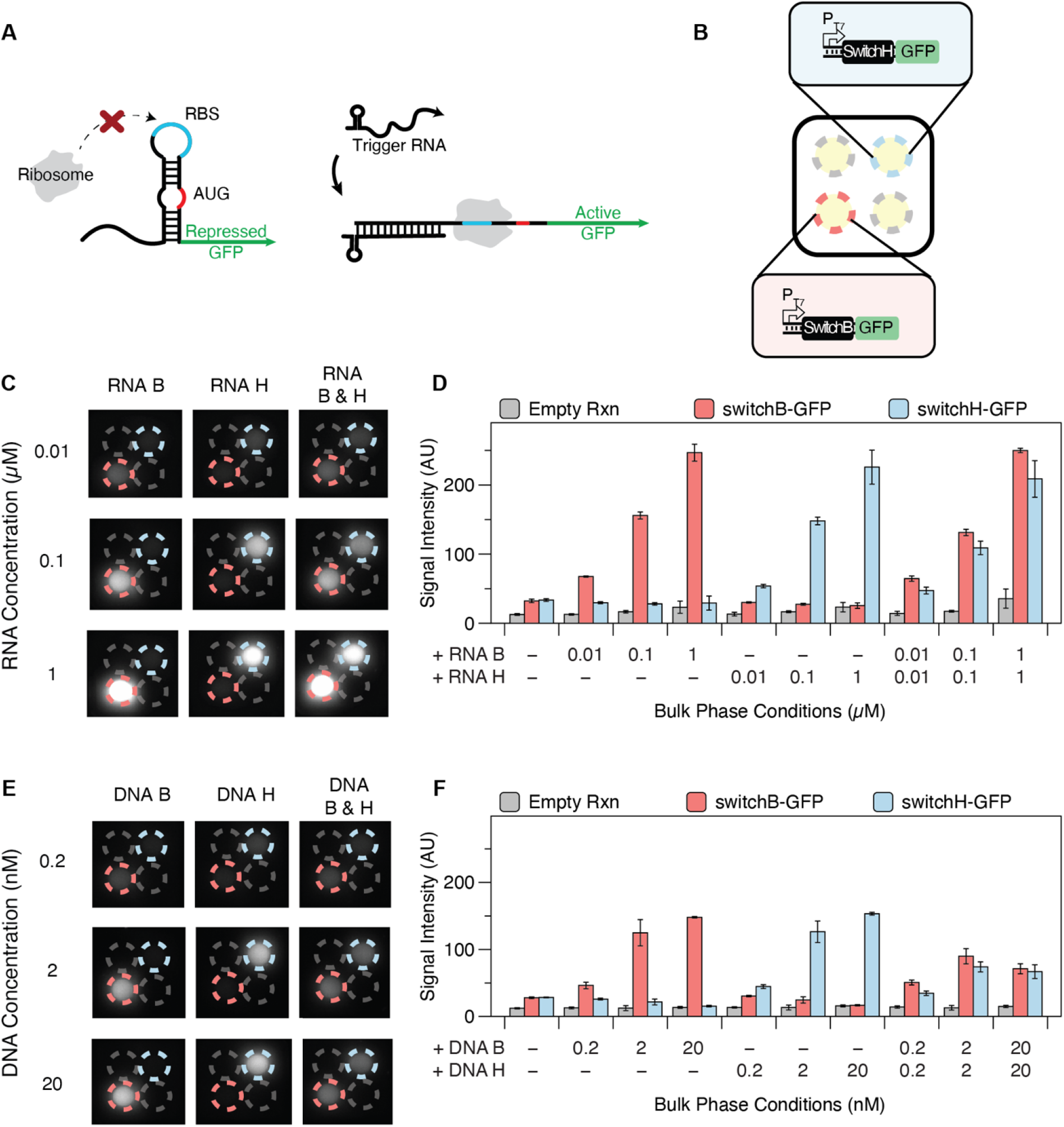
Multiplexed detection of model nucleic acid sequences in membrane-less protocell arrays. **(A)** Schematic of toehold switch mechanism. In the absence of trigger RNA, the switch RNA forms an inhibitory hairpin that blocks translation of a reporter (here, GFP). When trigger RNA is present, the switch RNA hybridizes with trigger RNA and unwinds the inhibitory hairpin, allowing translation of GFP. **(B)** Schematic of protocell array setup for multiplexed detection of nucleic acids. Red and blue circles indicate micro-basins containing toehold switches B and H, respectively. Gray circles indicate no-plasmid control protocells. **(C)** Representative fluorescence image of multiplexed RNA detection with minimal crosstalk. Images in the same column have the same RNA trigger(s) added. Images in the same row have the same concentration of trigger(s) added. Circle colors are as in (B). **(D)** Quantification of fluorescence images in (C) and their replicates. Error bars represent standard deviations of technical triplicates. **(E)** Representative fluorescence image of multiplexed detection of linear DNA with minimal crosstalk. Images in the same column have the same DNA trigger(s) added. Images in the same row have the same concentration of DNA trigger(s) added. Circle colors are as in (B). Although addition of both DNA triggers mutually represses their outputs, this was found to be specific to the use of these two triggers. **(F)** Quantification of fluorescence images in (E) and their replicates. Error bars represent standard deviations of technical triplicates.

Since DNA is more stable than RNA and can serve as a template for producing many RNA molecules, we hypothesized that protocell sensors could respond more sensitively to triggers expressed from a DNA template rather than direct detection of RNA. Furthermore, because RNA can be readily converted into DNA using reverse-transcriptase mediated isothermal amplification techniques like Nucleic Acid Sequence-Based Amplification (NASBA) or Recombinase Polymerase Amplification (RPA)^26,27^, sensing of DNA-templated RNA is a viable strategy for field-deployable detection of RNA targets. Just 2 nM of linear DNA (expressing RNA under control of a T7 promoter) added to the bulk phase strongly activated GFP expression, with slight activation detectable upon addition of just 0.2 nM of linear DNA (**Figures 4E, F**). Multiplexing using protocell arrays does not compromise sensitivity compared to CFE reactions (**Figure S4**). While the simultaneous addition of both triggers B and H caused some suppression of both signals compared to either one alone (**Figures 4D, F**), this effect was also observed in CFE reactions and was found to be specific only to triggers B and H (**Figure S5**), suggesting unfavorable sequence interactions independent of protocell array-based sensing.

### Multi-modal Detection of Clinically Relevant Targets in a Human Serum Matrix

Having demonstrated the multiplexing capabilities of protocell arrays, we next used this platform to detect multiple types of clinically relevant biomarker molecules with public health relevance: ions/minerals, small molecules, RNA, and DNA. The ions and small molecules we chose for detection were two micronutrients, zinc and vitamin B_12_ (adenosyl-cobalamin), that are important sensing targets for global health applications and for which our group has previous experience developing CFE biosensors^6,28^. The zinc sensor uses the activator ZntR, which activates expression from its cognate promoter P_ZntA_ when zinc is present. The B_12_ sensor uses the activator EutR, which activates expression from its cognate promoter P_EutS_ when both B_12_ and the cofactor ethanolamine (EA) are present. We characterized these micronutrient biosensors together in protocell arrays at concentrations near the physiologically relevant range to confirm sensor performance (**Figure S6**).

For nucleic acid detection, we developed and used toehold switch sensors that detect nucleic acid sequences from two bacterial pathogens: Shiga toxin-producing *E. coli* (STEC), which is a foodborne pathogen that causes life-threatening gastrointestinal symptoms^29^, and *Bacteroides thetaiotaomicron* (*B. theta*), which is linked to increased virulence of STEC^30^. We used a previously developed toehold switch for *B. theta*^31^, but had to design new switches for STEC, as previously published *E. coli* switches would cross-react with non-pathogenic *E. coli* strains^31^. We developed two STEC switches targeting genomic sequences of toxin proteins: Shiga toxin I (Stx1) and Shiga toxin II (Stx2)^32^. When different combinations of linear DNA coding for *B. theta*, Stx1, and Stx2 triggers were added to the bulk phase of a microwell containing protocell arrays, all protocell sensors showed orthogonal trigger detection with minimal reaction crosstalk (**Figure S7**), demonstrating that protocell arrays can reliably multiplex the detection of pathogenic bacteria nucleic acids.

We then combined the validated micronutrient and bacterial sensors to demonstrate multi-modal, multiplexed detection of diverse classes of analytes from a chemically defined sample and from a contrived human serum sample. A protocell array containing sensors for zinc, B_12_, Stx1, and Stx2 was added to the microwell along with different combinations of analytes (zinc, B_12_, Stx1 trigger RNA, and/or linear DNA for Stx2 trigger expression) (**Figure 5A**). Each protocell sensor produced GFP only when its cognate analyte was present, demonstrating successful multiplexed detection spanning multiple molecular classes (ion, small molecule, RNA, and DNA) from a single sample (**Figure 5B**). With this multi-modal multiplexed sensing platform established, we next sought to verify that sensing with membrane-less protocell arrays remains robust even in complex samples such as human serum. We spiked 20% human serum (similar to sample concentrations used in our previously published CFE efforts)^6^ into the bulk phase and observed the same target-specific activation from all protocell sensors (**Figure 5C**), demonstrating our platform’s translational potential for assessment of biomarkers in patient samples. We note that the presence of Stx1 RNA and B_12_ caused a slight increase in overall (background and activated) expression from the other two sensors. This is a prototypical example of a “matrix effect”, a widely-encountered analytical issue that must be addressed for almost any quantitative sensor or diagnostic^6,33–35^.

**Figure 5:**
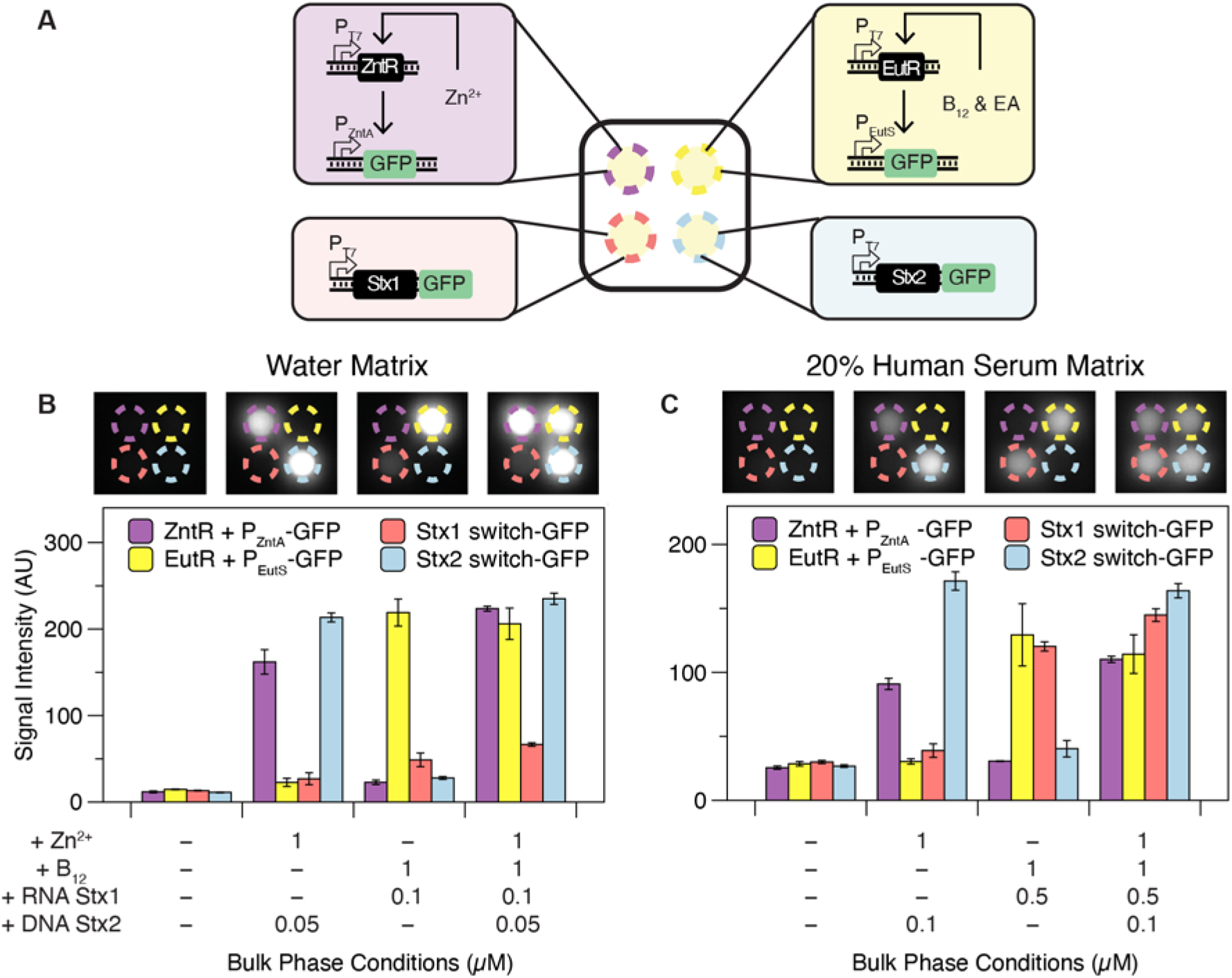
Multiplexed detection of clinically relevant biomarkers across molecular classes in water and 20% human serum matrices. (**A**) Schematic of membrane-less protocell array setup for multi-modal, multiplexed detection of diverse classes of clinically relevant biomarkers. Purple and yellow circles indicate micro-basins containing zinc and vitamin B_12_ sensors, respectively. Red and blue circles indicate micro-basins containing Stx1 and Stx2 toehold switches, respectively. (**B**) Representative fluorescence image (top) and quantification of fluorescence of representative image and its replicates (bottom) demonstrating multi-modal target measurement multiplexing using protocell arrays, with simultaneous detection of ion, small molecule, RNA, and DNA targets from the same bulk phase. Error bars represent standard deviations of technical triplicates. (**C**) Multi-modal target detection in a 20% human serum matrix. Representative fluorescence image (top) and quantification of fluorescence of representative image and its replicates (bottom) demonstrating robustness of protocell arrays to a complex sample matrix. Error bars represent standard deviations of technical triplicates.

### Toward a Field-Deployable, Equipment-Free Diagnostic Platform

To be useful as a minimal-equipment, point-of-care sensing or diagnostic tool, the membrane-less protocell array platform should produce test results that can be visually interpreted without the aid of equipment like plate readers or fluorescence imagers. Toward this goal, we replaced the GFP reporter protein with β-galactosidase (LacZ), an enzyme that catalyzes the production of a colorimetric readout. We tested two colorimetric substrates, chlorophenol red-beta-D-galactopyranoside (CPRG) and 5-Bromo-4-chloro-3-indolyl β-D-galactopyranoside (X-gal), that are substrates for LacZ and are widely used for cell-free diagnostics and molecular biology assays, respectively^6,7,24,31,36^. We also used a newly-prepared white microwell plate to enable improved pigment visualization, reduced the bulk phase volume by half, and increased the number of micro-basins in a standard 96-well plate microwell to multiplex more reactions (**Figure S1B**). We found that while LacZ can produce visible color change faster when cleaving CPRG than when cleaving X-gal, the product of CPRG cleavage rapidly diffuses from the protocell to the surrounding bulk phase (**Figure S8**). This diffusion could potentially obscure result interpretation if readings are not to be taken within 30 minutes of color change, which could be an issue if different multiplexed sensors have different characteristic response times. As a result, we chose to use X-gal for subsequent test development based on its stable localization of pigments, despite its longer reaction time.

We next used this colorimetric readout for detection of the presence of nucleic acid sequences characteristic of pathogenic bacteria. A protocell array with toehold switch-based sensors for *B. theta*, Stx1, and Stx2 was loaded into micro-basins, and the bulk phase contained combinations of linear DNA coding for expression of cognate triggers (**Figure 6A)**. Within three hours, all protocell sensors turned visibly blue to accurately report on the presence of the triggers in the bulk phase, with minimal pigment leakage or sensor crosstalk (**Figure 6B**).

**Figure 6:**
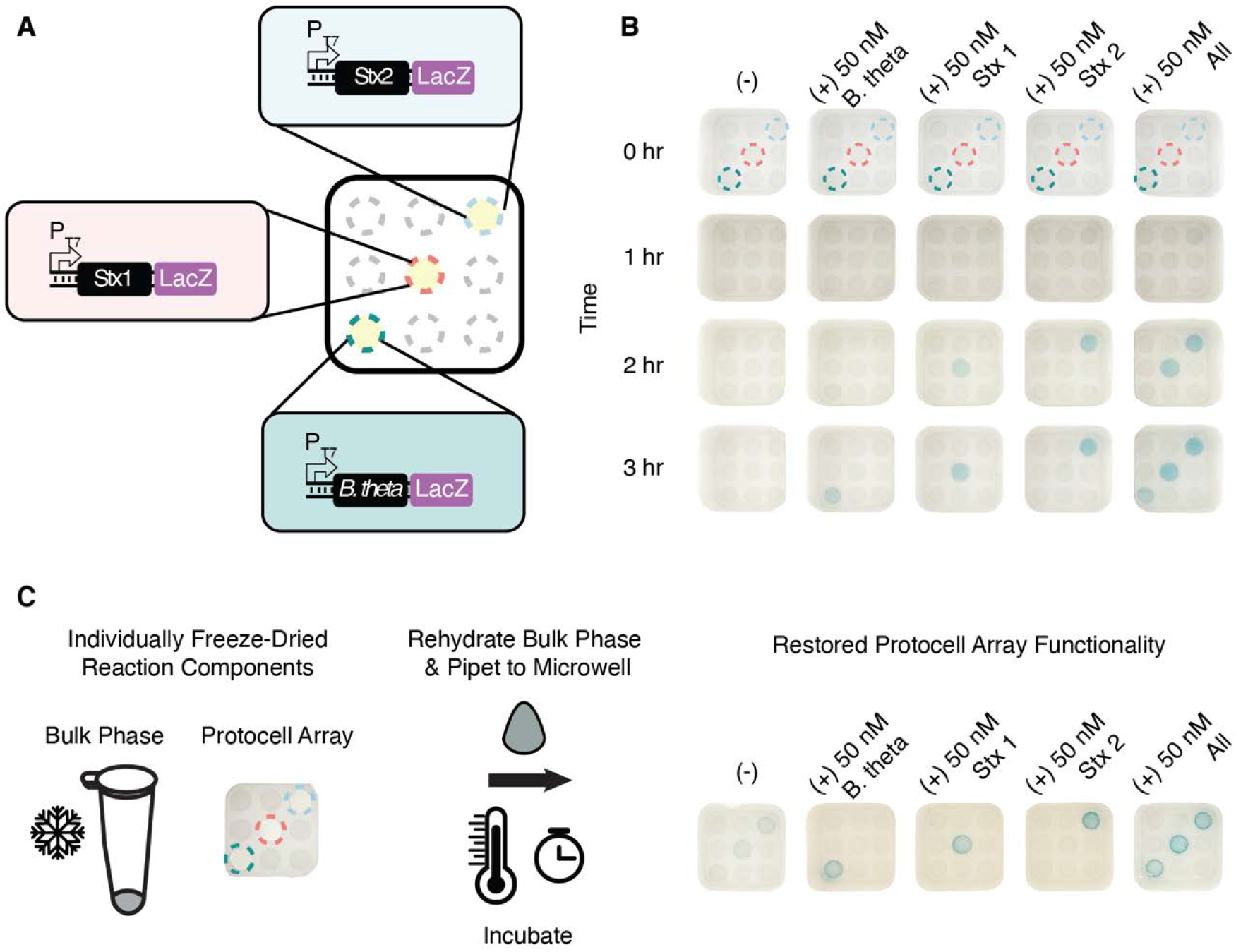
Membrane-less protocell arrays meet key criteria for minimal equipment, field-deployable multiplexed assays. (**A**) Schematic of colorimetric protocell array setup for multiplexed detection of nucleic acids. Teal, red, and blue circles indicate micro-basins containing *B. theta*, Stx1, and Stx2 protocell sensors, respectively. Gray circles indicate empty micro-basins without CFE reactions. (**B**) Representative sensor activation and pigment production in different bulk phase conditions and at different time points. Images in the same column have the same RNA trigger(s) added. Images in the same row were taken after the same incubation time. Dashed circles indicate micro-basins containing protocell sensors as shown in (A). (**C**) Protocell array functionality after lyophilization. Protocells containing CFE reactions coding for sensors of bacterial nucleic acid sequences were lyophilized in micro-basins within microwells, while bulk phase solutions were lyophilized separately. Addition of sample-reconstituted bulk phase to microwells caused rehydration of the dehydrated membrane-less protocells, reformed liquid-liquid phase separation, and revived CFE sensing reactions leading to pigment production.

Additionally, a field-deployable protocell array-based sensing platform must be compatible with storage and transportation at ambient temperature for easy distribution to the point of need. To demonstrate our system’s ability to meet this criterion, we deposited protocells containing sensors for different nucleic acid sequences in the multiplexing micro-basins and lyophilized them. We separately lyophilized a bulk phase solution containing PEG, X-gal, and cell-free energy mix. Addition of water containing cognate targets for the sensors to the lyophilized PEG solution reconstituted the bulk phase, which was then added to the microwells. This rehydrated the protocell array and re-formed the liquid-liquid phase separation. The CFE sensing reactions were also revived while remaining compartmentalized inside their protocells and producing visible color readouts after incubation (**Figure 6C**). This result suggests the possibility for a simple-to-operate multiplexing assay that can be performed by minimally-trained staff, which is critical for point-of-need use in the field in, for example, environmental or epidemiological surveillance.

## Discussion

Membrane-less protocell models have in recent years been used for a variety of scientific and engineering applications, including cell-like functionalities such as selective nucleic acid retention^15,37^, reaction acceleration^38,39^, and droplet division^40^. Despite being membrane-less, these phase-separated droplets can be quite robust, maintaining their structure through dehydration and rehydration cycles^41,42^ and enabling potentially impactful downstream uses. One example is the acoustically-trapped protocell patterning technique that enables different enzymatic reactions to occur in adjacent protocells^43^. However, such a strategy requires complex equipment and extensive optimization of reaction environments, making it difficult to implement at the point of need.

Our approach using polymer ATPS to construct protocell arrays can address current limitations in bringing multiplexed analyte measurement to field applications. We show that topographical micro-basin features patterned on the floor of standardized 96-well plate format microwells (**Figure S1**) enables multiple protocells to coexist spatially separated with minimal sensor interference and without requiring any external input to maintain protocell separation. The membrane-less aspect of protocells formed by ATPS also facilitates minimally-hindered analyte diffusion (and even concentration, for some analytes with favorable partitioning) into the reaction droplets. Furthermore, these protocells compartmentalize custom-designed CFE reactions that can quantify diverse classes of analytes, remain robust to complex sample environments, and retain their sensing capabilities after lyophilization for on-demand multiplexed analyte measurement at the point of need.

The embedding of CFE reactions in ATPS-formed protocells is uniquely poised to meet multiple key challenges in biosensing applications, as this platform can be customized to detect diverse sets of analytes merely by reprogramming the DNA sequence of plasmids used as the basis for their sensing. Previous work has demonstrated that CFE can be used to detect metabolites *via* incorporation of genetically encoded metabolic transducers^11,44^, proteins *via* genetically encoded aptamers^45,46^, and nucleic acids *via* genetically encoded synthetic regulators^7,8,31^. These different sensing systems may require differently optimized CFE reactions (e.g., lysate preparations or macromolecular reaction additives), but these individually optimized sub-systems can all be used together in parallel to measure different targets in the same sample; of note, the protocells sensing nucleic acids in **Figure 5** use different lysate preparations than the other protocells in **Figure 5**. Being able to selectively tune an individual sensor’s performance without negatively impacting other sensors is an enabling advance for multiplexed measurement of analytes, made possible by encapsulation of individual sensors into discrete, phase-separated protocells. Furthermore, compartmentalization of protocell sensors eliminates the need for multiple, orthogonal, or target-specific reporter systems, allowing all sensors in the same microwell to be developed with the same reporter system. This facilitates rapid platform development, as newly developed sensors can be readily incorporated into existing protocell arrays with obsolete sensors easily replaced.

An array of spatially separated gene networks that sense and respond to different analytes in the same environment has many potential applications. One such example is in agricultural and environmental surveillance^3^, where a small environmental sample can indicate the presence of a pollutant, excessive fertilizer, or even an animal or plant pathogen. Protocell arrays could also be used as a platform for prototyping chemical communication networks among synthetic cells^47,48^. Each protocell can be separately programmed with a different function, allowing for investigation of communication at the single-(proto)cell level that could allow characterization of phenomena including stochasticity in these communication networks.

Another potential application of these membrane-less protocell arrays is for point-of-care diagnostics with multiplexed detection of clinically relevant biomarkers in human biofluid sample matrices. Many diseases and disorders that clinicians and researchers seek to identify in the field are not diagnosed based on just one biomarker, but by the combination of multiple test results often spanning classes of biomolecules. The use of protocell arrays formed by CFE and ATPS is an enabling platform for these needs. It is compatible with complex sample matrices and can withstand lyophilization for intact functionality upon rehydration, enabling flexible test development and deployment to the point of need without expensive cold chain requirements. However, in our efforts to demonstrate serum compatibility with a protocell array platform, we found that serum proteins formed cloudy precipitates when added to the PEG-rich bulk phase. While serum protein precipitation did not obstruct biphasic polymer separation or detection of GFP signal, it hindered pigment visualization in colorimetric reactions (data not shown). However, there are clear paths to mitigating this shortcoming, thereby enabling multiplexed serum biomarker detection in a minimal-equipment format.

To be effectively used for the detection of pathogenic or host nucleic acids, protocell arrays should be integrated with nucleic acid amplification steps. Most nucleic acid targets are present at attomolar to femtomolar levels in biological samples, which is below the detection limit of cell-free nucleic acid sensors^7,9,31^. It has previously been shown that nucleic acids can be amplified in the field through isothermal processes (e.g., NASBA or RPA)^7–9,26,27,31^, but these steps admittedly increase the level of complexity of the test and require trained operators to execute. Although nearly all current CFE nucleic acid sensors require an amplification step to detect relevant concentrations of sequences, there are numerous ongoing research efforts aiming to eliminate the need for upstream input amplification. Given the demonstrated modularity and easily reconfigurable format of our platform, those resulting amplification-free CFE systems could be easily incorporated into our protocell array and enable one-pot detection of clinically relevant biomarker levels. Nonetheless, one would still need to ensure that any upstream processing does not alter levels of other targeted biomarkers present in the sample.

Other future advancements could come from further miniaturization to reduce required sample volumes. While the final volume of 11 μL per each of 9 assays is consistent with previous CFE-based sensors^4–8^, there may be applications where 100 μL of sample is not always accessible. There may be lower limits on the total sample volume based on the requirement for visibility of multiplexed readouts to the naked eye, but use of a smartphone app for image acquisition and data processing could supplement completely visual detection.

In conclusion, we have demonstrated that arrays of polymer ATPS-formed, membrane-less protocells with compartmentalized DNA transcription and RNA translation machinery can perform multiplexed detection of diverse classes of analytes. Such a protocell system survives lyophilization to enable test storage and distribution at ambient temperature. Rehydration with an analyte-containing polymer solution reestablishes phase separation, allows uptake of analytes by the resulting protocell sensors, and revives compartmentalized transcription- and translation-mediated detection reactions. These arrays of membrane-less protocells provide modular, field-deployable, multiplexed diagnostic potential. Interfacing CFE reactions with membrane-less protocells addresses the current limitation in multiplexed detection of diverse analyte from a single sample and opens new opportunities for implementing diagnostic panels for use at the point of care.

## Methods

### Bacterial Strains

*E. coli* strain DH10β was used for all cloning and plasmid preparations. *E. coli* strain BL21 Star (DE3) ∆*lacIZYA* was created by lambda red recombination^49^ and used for in-house cell-free lysate preparation. Genomic DNA from *E. coli* strains DH10β and BL21 Star (DE3) were used as negative controls for target-specific amplification of Stx1 and Stx2 triggers. Genomic DNA from *B. thetaiotaomicron* (ATCC 29148D) and *E. coli* O157:H7 (ATCC 51657GFP) were used for detection of pathogenic bacteria.

### Genetic Parts Assembly and Plasmid Preparation

Supplementary **Table S1** contains sequences of all parts used in this study. DNA oligonucleotides for cloning and sequencing were synthesized by Eurofins Genomics. Partial sequences for small molecule sensors and toehold switches were obtained from previously published sequences and were synthesized either as gene fragments or ssDNA-annealed oligonucleotides from Eurofins Genomics. Plasmids expressing regulators and reporter proteins were cloned using either Gibson Assembly^50^ or blunt end ligation into plasmid backbone pJL1. Assembled constructs were transformed into DH10β cells, and isolated colonies were grown overnight in LB with antibiotics. Plasmid DNA from overnight cultures was purified using EZNA miniprep columns (OMEGA Bio-Tek). Plasmid sequences were verified with Sanger DNA sequencing (Eurofins Genomics).

Plasmid DNA used for all cell-free and protocell reactions was purified from EZNA midiprep columns (OMEGA Bio-Tek) followed by isopropanol and ethanol precipitation. The purified DNA pellet was reconstituted in elution buffer, measured on a Nanodrop 2000 for concentration, and stored at −20 °C.

Genomic DNA for *B. thetaiotaomicron* (ATCC 29148D) used for pathogenic detection was purchased from American Type Culture Collection (ATCC). Genomic DNA for *E. coli* O157:H7 (ATCC 51657GFP) used for pathogenic detection was obtained from overnight culture grown in tryptic soy broth supplemented with 1% glucose at 37 °C. DNA was extracted using an Invitrogen PureLink Microbiome DNA Purification Kit (A29790). Genomic DNA for DH10β and BL21 Star (DE3) was obtained from a 5 mL overnight culture grown in LB medium. DNA was extracted using Quick-DNA Plus Kit (Zymo Research) according to the manufacturer’s protocol.

### Preparation of In-House Cell-Free Lysate

Cellular lysate for all experiments is prepared as described by Sun *et al* ^51^ with a few protocol modifications. Briefly, BL21 Star (DE3) ∆*lacIZYA* cells were grown in 2× YTP medium at 37 °C and 220 rpm to an optical density (OD) between 1.5-2.0, corresponding to the mid-exponential growth phase. Lysate prepared for toehold switch expression had an additional IPTG (0.4 mM) induction step when the OD reached 0.4 to activate expression of T7 RNA polymerase, creating a T7 RNAP-enriched lysate. Cells were centrifuged at 2700 rcf and washed three times with S30A buffer (50 mM tris, 14 mM magnesium glutamate, 60 mM potassium glutamate, 2 mM dithiothreitol, and pH-corrected to 7.7 with acetic acid). Cells were then centrifuged at 2700 rcf and washed three times with S30 buffer. After the final centrifugation, the wet cell mass was determined, and cells were resuspended in 1 mL of S30A buffer per 1 g of wet cell mass. The cellular resuspension was divided into 1 mL aliquots. Cells were lysed using a Q125 sonicator (Qsonica) at a frequency of 20 kHz and at 50% of amplitude. Cells were sonicated on ice with cycles of 10 s on and 10 s off, delivering approximately 200 J, at which point the cells appeared visibly lysed. An additional 4 mM dithiothreitol was added to each tube, and the sonicated mixture was then centrifuged at 12,000 rcf and 4°C for 10 min. After centrifugation, the supernatant was removed, divided into 0.5 mL aliquots, and incubated at 37°C and 220 rpm for 80 min. After this runoff reaction, the cellular lysate was centrifuged at 12,000 rcf and 4°C for 10 min. The supernatant was removed and loaded into a 10 kDa molecular weight cutoff dialysis cassette (Thermo Fisher). Lysate was dialyzed in 1 liter of S30B buffer (14 mM magnesium glutamate, 60 mM potassium glutamate, 1 mM dithiothreitol, and pH-corrected to 8.2 with tris) at 4°C for 3 hours. Dialyzed lysate was removed and centrifuged at 12,000 rcf and 4°C for 10 min. The supernatant was removed, aliquoted, flash-frozen in liquid nitrogen, and stored at −80°C for future use.

### Cell-Free Reactions

Cell-free reaction were assembled as previously described by Kwon and Jewett ^52^. Briefly, reaction mixtures were composed of 27 v/v% of in-house prepared lysate, 2 mM each proteinogenic amino acid, 1.2 mM ATP, 0.85 mM each of GTP, CTP, and UTP, 0.2 mg/mL tRNA, 0.27 mM CoA, 0.33 mM NAD, 0.068 mM folinic acid, 1.5 mM spermidine, 33 mM PEP, 130 mM potassium glutamate, 10 mM Ammonium glutamate, 12 mM magnesium glutamate, 4 mM sodium oxalate, and specified concentrations of plasmids (described in Table S2), RNA triggers, and small molecules. Each assembled cell-free reaction was 10 μL in volume and placed in a black-bottomed 384-well plate (Greiner Bio-One) and incubated at 37 °C for 3 hours for GFP expression. A clear adhesive film was used to cover the plate and prevent evaporation.

### Protocell CFE Reactions

Polymers used to establish ATPS-based membrane-less protocells were prepared by dissolving either 400k Ficoll, 500k dextran, or 35k PEG into nuclease-free water. The bulk phase at time of preparation and before adding to the Ficoll or dextran phase solution consists of 5% (v/v) of 35k PEG, 1x concentration of all reagents added for cell-free reactions (excluding lysate), specified concentrations of small molecules or nucleic acids, and nuclease-free water to a final volume of 200 μL for the 4-plex system or 100 μL for the 9-plex system. For experiments with human serum, RNase Inhibitor, Murine, (New England Biolabs) was added to a concentration of 0.6 U/μL in the bulk phase. For colorimetric output in cell-free reactions, color substrates were also added to the bulk phase to a final concentration of 0.6 mg/mL for CPRG or 0.2 mg/mL for X-gal.

Concentrations for individual sensor reactions used in each figure are compiled in **Table S2**. Briefly, each protocell sensor consisted of 10% (v/v) Ficoll or dextran polymers at time of preparation, 27% (v/v) cell-free lysate, 1x concentration of cell-free reagents, specific concentrations of plasmid DNA, and water to a final volume of 2 μL for the 4-plex system or 1 μL for the 9-plex system. For detection of linear DNA added in the bulk phase, each protocell contained either 2 μM of χDNA or 2 μM of the GamS protein (Arbor Bioscience) for protection against endonucleases present in cell-free lysate. The assembled protocell solution was then vortexed at a medium-high setting to ensure homogenous mixing.

To assemble the protocell arrays, bulk phase solution containing specified concentrations of targets was first pipetted into the custom plate (PHASIQ) to fill up each microwell. Protocell droplets were then pipetted into the bulk phase solution at designated micro-basin. Unless otherwise noted, microwells containing assembled protocell arrays were all incubated at 37 °C for 3 hours before imaging on a ChemiDoc MP (Bio-Rad) imaging system. A clear adhesive film was used to cover the plate and prevent evaporation.

### Image Acquisition and Data Analysis

For cell-free reactions, end point GFP readings were taken using a plate reader (Synergy4, BioTek). The excitation and emission wavelengths were 485 and 528 nm, with a gain setting of 70. For protocell array reactions with fluorescent reporters, the ChemiDoc MP imaging system (Bio-Rad) was used for fluorescent plate imaging. Image Lab software (Bio-Rad) was used for image collection with settings of 0.5 s exposure time, grayscale image color, 530/28 filter for GFP detection, and Blue Epi illumination as a light source. An image transform procedure was uniformly applied to all images in Image Lab (with high, low, and gamma values of 10000, 0, and 1, respectively) before exporting files for analysis. For image analysis, each image was first converted to an 8-bit grayscale image. An image processing software, Fiji, was used to extract signal intensities from each 1.5 mm diameter micro-well (pixel size 50) for data analysis. For colorimetric protocell array reactions, all pictures were taken with an iPhone X (Apple) in a light-controlled setting^53^. Adobe Photoshop was used to crop individual wells for data presentation. A brightness filter was uniformly applied to all colorimetric ATPS photos to make them better resemble appearance to the naked eye.

### Trigger Preparation

DNA encoding each trigger RNA used in experiments was either amplified from cloned plasmid or from genomic DNA of specified species of bacteria by PCR with Q5 DNA polymerase (New England Biolabs). Primers were designed to create linear DNA with a T7 promoter as well as additional protective sequences on the 5’ and 3’ ends of linear template. Sequences for primers used to amplify triggers from DNA template or genomic DNA are provided in supplementary **Table S3**. After PCR amplification, all products were run on a 2 w/v% agarose gel to verify successful amplification of target and then purified using a PCR purification kit (Omega Bio-Tek). The prepared linear DNA was either directly used in cell-free and cell-free ATPS reactions or used as a template for *in vitro* transcription.

RNA triggers were transcribed from linear DNA template using T7 polymerase according to the manufacturer’s protocol (New England Biolabs). Following RNA synthesis, Dnase I (Zymo Research) was added to degrade linear DNA template. The RNA products were then purified using an RNA Clean and Concentrator kit (Zymo Research) according to the manufacturer’s protocol. Following purification, RNA concentration was measured on a Nanodrop 2000, subaliquoted, and stored −20 °C.

### STEC Toehold Switch Development

Toehold switches targeting Shiga toxins in STEC (Stx1 and Stx2) were designed using NUPACK with series B toehold switch design^25^ and cloned into a pJL1 plasmid containing a GFP reporter. Trigger sequences were synthesized by Eurofins Genomics as gene fragments containing a T7 promoter and 30-35 bp of protective sequences before and after the actual trigger sequence and cloned onto the pJL1 plasmid backbone to facilitate sensor screening. Trigger/sensor pairs were then tested in cell-free reactions containing 2.5 nM toehold switch sensor-GFP reporter plasmid and 5 nM of trigger plasmid to verify successful sensor activation and orthogonality (**Figure S9**). Following trigger/sensor pair validation, primers used to amplify different trigger sequences from genomic DNA were verified for specificity toward their targets using PCR reactions with non-target templates (**Figure S10**).

### Serum Processing

Pooled human serum was purchased from Corning (Corning, NY). Endogenous zinc was removed from serum through Chelex-100 treatment. In total, 1 g of Chelex-100 resin was added to 100 mL of serum, and the mixture was vigorously stirred for 2 hrs at room temperature. Resin was then isolated from the serum through centrifugation followed by syringe filtering. All serum samples were aliquoted to minimize freeze-thaw cycles and stored at −20 °C until use.

Measurement of successful zinc removal from serum was conducted at the University of Georgia Laboratory for Environmental Analysis (**Figure S11**). Samples were digested with concentrated acid and analyzed on ICP-MS according to EPA method 3052.

### Lyophilization

Protocell sensors containing 55% (v/v) lysate, specified concentrations of plasmid DNA (**Table S2**), and 10% (v/v) Ficoll were directly added to micro-basins and the plate was stored at −80 °C overnight to freeze. Bulk phase solutions containing PEG, 0.2 mg/mL X-gal, and cell-free reagents at 1 x concentration were prepared in 1.5 mL Eppendorf tubes and stored at −80 °C overnight to freeze. Frozen plates and bulk phase solutions were removed the next day and added to a pre-chilled Labconco Fast-Freeze flask. Samples were transferred as quickly as possible to prevent reagent thawing. Flasks were connected to a Labconco benchtop lyophilizer and lyophilized at −50 °C and 0.05 mbar for 3 hours.

100 μL of water mixed with 50 nM of specified bacterial triggers was used to reconstitute the lyophilized bulk phase solution. Reconstituted bulk phase solutions were added to microwells at room temperature before incubating at 37 °C. A clear adhesive film was used to cover the plate and prevent evaporation.

## Acknowledgement

We thank Dr. Michael Jewett for his gift of the pJL1 plasmid. We thank R. Kane and his laboratory for usage of and assistance with their ChemiDoc MP imaging system. We thank P. Santangelo and his laboratory for usage of and assistance with their lyophilizer.

## Author Contributions

Conceptualization: YZ, TK, MPM, MPS and ST; Investigation: YZ, TK, GK, and MPM; Formal Analysis: YZ, TK; Writing – Original Draft: YZ; Writing – Review & Editing: YZ, MPM, TK, GK, MPS, ST; Visualization: YZ; Supervision: MPS and ST; Funding Acquisition: MPS and ST.

