## Supplementary Tables for "Protocell Arrays for Simultaneous Detection of Diverse Analytes"

**Table S1:** Description of plasmid parts and DNA sequences in this paper.

<table>
<tr><th>Name</th><th>Construct Description</th></tr>
<tr><td>pJL1</td><td>Plasmid encoding superfolder GFP expression under P<sub>T7</sub> promoter with strong ribosomal binding site</td></tr>
<tr><td colspan="2">P<sub>T7</sub> Promoter-Stability Hairpin-StrongRBS-sfGFP-T7 Terminator<br><br>taatacgactcactatagggagaccacaacggtttccctctagaaataattttgtttaactttaagaaggagatatacatATGAGC<br>AAAGGTGAAGAACTGTTTACCGGCGTTGTGCCGATTCTGGTGGAACTGGATGGCGATGT<br>GAACGGTCACAAATTCAGCGTGCGTGGTGAAGGTGAAGGCGATGCCACGATTGGCAAA<br>CTGACGCTGAAATTTATCTGCACCACCGGCAAACTGCCGGTGCCGTGGCCGACGCTGG<br>TGACCACCCTGACCTATGGCGTTCAGTGTTTTAGTCGCTATCCGGATCACATGAAACGTC<br>ACGATTTCTTTAAATCTGCAATGCCGGAAGGCTATGTGCAGGAACGTACGATTAGCTTTA<br>AAGATGATGGCAAATATAAAACGCGCGCCGTTGTGAAATTTGAAGGCGATACCCTGGTG<br>AACCGCATTGAACTGAAAGGCACGGATTTTAAAGAAGATGGCAATATCCTGGGCCATAAA<br>CTGGAATACAACTTTAATAGCCATAATGTTTATATTACGGCGGATAAACAGAAAAATGGCA<br>TCAAAGCGAATTTTACCGTTCGCCATAACGTTGAAGATGGCAGTGTGCAGCTGGCAGAT<br>CATTATCAGCAGAATACCCCGATTGGTGATGGTCCGGTGCTGCTGCCGGATAATCATTAT<br>CTGAGCACGCAGACCGTTCTGTCTAAAGATCCGAACGAAAAAGGCACGCGGGACCACAT<br>GGTTCTGCACGAATATGTGAATGCGGCAGGTATTACGTGGAGCCATCCGCAGTTCGAAA<br>AATAAgtcgaccggctgctaacaaagcccgaaaggaagctgagttggctgctgccaccgctgagcaataactagcataaccc<br>cttggggcctctaaacgggtcttgaggggttttttg</td></tr>
</table>

<table>
<tr><td>P<sub>T7</sub>-LacI</td><td>Plasmid encoding LacI expression under P<sub>T7</sub> promoter with strong ribosomal binding site</td></tr>
<tr><td colspan="2">P<sub>T7</sub> Promoter-Stability Hairpin-StrongRBS-LacI-T7 Terminator<br><br>taatacgactcactatagggagaccacaacggtttccctctagaaataattttgtttaactttaagaaggagatatacatATGAAA<br>CCAGTAACGTTATACGATGTCGCAGAGTATGCCGGTGTCTCTTATCAGACCGTTTCCCG<br>CGTGGTGAACCAGGCCAGCCACGTTTCTGCGAAAACGCGGGAAAAAGTGGAAGCGGCG<br>ATGGCGGAGCTGAATTACATTCCCAACCGCGTGGCACAACAACTGGCGGGCAAACAGT<br>CGTTGCTGATTGGCGTTGCCACCTCCAGTCTGGCCCTGCACGCGCCGTCGCAAATTGTC<br>GCGGCGATTAAATCTCGCGCCGATCAACTGGGTGCCAGCGTGGTGGTGTCGATGGTAG<br>AACGAAGCGGCGTCGAAGCCTGTAAAGCGGCGGTGCACAATCTTCTCGCGCAACGCGT<br>CAGTGGGCTGATCATTAACTATCCGCTGGATGACCAGGATGCCATTGCTGTGGAAGCTG<br>CCTGCACTAATGTTCCGGCGTTATTTCTTGATGTCTCTGACCAGACACCCATCAACAGTA<br>TTATTTTCTCCCATGAAGACGGTACGCGACTGGGCGTGGAGCATCTGGTCGCATTGGGT<br>CACCAGCAAATCGCGCTGTTAGCGGGCCCATTAAGTTCTGTCTCGGCGCGTCTGCGTCT<br>GGCTGGCTGGCATAAATATCTCACTCGCAATCAAATTCAGCCGATAGCGGAACGGGAAG<br>GCGACTGGAGTGCCATGTCCGGTTTTCAACAAACCATGCAAATGCTGAATGAGGGCATC<br>GTTCCCACTGCGATGCTGGTTGCCAACGATCAGATGGCGCTGGGCGCAATGCGCGCCA<br>TTACCGAGTCCGGGCTGCGCGTTGGTGCGGATATCTCGGTAGTGGGATACGACGATAC<br>CGAAGACAGCTCATGTTATATCCCGCCGTTAACCACCATCAAACAGGATTTTCGCCTGCT<br>GGGGCAAACCAGCGTGGACCGCTTGCTGCAACTCTCTCAGGGCCAGGCGGTGAAGGG<br>CAATCAGCTGTTGCCCGTCTCACTGGTGAAAAGAAAAACCACCCTGGCGCCCAATACGC<br>AAACCGCCTCTCCCCGCGCGTTGGCCGATTCATTAATGCAGCTGGCACGACAGGTTTCC<br>CGACTGGAAAGCGGGCAGTGATAAgtcgaccggctgctaacaaagcccgaaaggaagctgagttggctgctgc<br>caccgctgagcaataactagcataaccccttggggcctctaaacgggtcttgaggggttttttg</td></tr>
</table>

<table>
<tr><td>P<sub>T7lacO</sub>-GFP</td><td>Plasmid encoding superfolder GFP expression under P<sub>T7lacO</sub> promoter with strong ribosomal binding site</td></tr>
<tr><td colspan="2">PT7 Promoter-lacO-Stability Hairpin-StrongRBS-sfGFP- T7 Terminator<br><br>taatacgactcactatagggagatgtgagcggataacaaccacaacggtttccctctagaaataattttgtttaactttaagaaggag<br>atatacatATGAGCAAAGGTGAAGAACTGTTTACCGGCGTTGTGCCGATTCTGGTGGAACTG<br>GATGGCGATGTGAACGGTCACAAATTCAGCGTGCGTGGTGAAGGTGAAGGCGATGCCA<br>CGATTGGCAAACTGACGCTGAAATTTATCTGCACCACCGGCAAACTGCCGGTGCCGTGG<br>CCGACGCTGGTGACCACCCTGACCTATGGCGTTCAGTGTTTTAGTCGCTATCCGGATCA<br>CATGAAACGTCACGATTTCTTTAAATCTGCAATGCCGGAAGGCTATGTGCAGGAACGTAC<br>GATTAGCTTTAAAGATGATGGCAAATATAAAACGCGCGCCGTTGTGAAATTTGAAGGCGA<br>TACCCTGGTGAACCGCATTGAACTGAAAGGCACGGATTTTAAAGAAGATGGCAATATCCT<br>GGGCCATAAACTGGAATACAACTTTAATAGCCATAATGTTTATATTACGGCGGATAAACA<br>GAAAAATGGCATCAAAGCGAATTTTACCGTTCGCCATAACGTTGAAGATGGCAGTGTGCA<br>GCTGGCAGATCATTATCAGCAGAATACCCCGATTGGTGATGGTCCGGTGCTGCTGCCGG<br>ATAATCATTATCTGAGCACGCAGACCGTTCTGTCTAAAGATCCGAACGAAAAAGGCACGC<br>GGGACCACATGGTTCTGCACGAATATGTGAATGCGGCAGGTATTACGTGGAGCCATCCG<br>CAGTTCGAAAAATAAgtcgaccggctgctaacaaagcccgaaaggaagctgagttggctgctgccaccgctgagcaat<br>aactagcataaccccttggggcctctaaacgggtcttgaggggttttttg</td></tr>
</table>

<table>
<tr><td>AraC-P<sub>BAD</sub>-GFP</td><td>Plasmid with divergent P<sub>BAD</sub> promoter for constitutive AraC expression and arabinose modulated superfolder GFP expression.</td></tr>
<tr><td colspan="2">AraC-PBAD-Stability Hairpin-StrongRBS-sfGFP- T7 Terminator<br><br>TTATGACAACTTGACGGCTACATCATTCACTTTTTCTTCACAACCGGCACGGAACTCGCT<br>CGGGCTGGCCCCGGTGCATTTTTTAAATACCCGCGAGAAATAGAGTTGATCGTCAAAAC<br>CAACATTGCGACCGACGGTGGCGATAGGCATCCGGGTGGTGCTCAAAAGCAGCTTCGC<br>CTGGCTGATACGTTGGTCCTCGCGCCAGCTTAAGACGCTAATCCCTAACTGCTGGCGGA<br>AAAGATGTGACAGACGCGACGGCGACAAGCAAACATGCTGTGCGACGCTGGCGATATC<br>AAAATTGCTGTCTGCCAGGTGATCGCTGATGTACTGACAAGCCTCGCGTACCCGATTAT<br>CCATCGGTGGATGGAGCGACTCGTTAATCGCTTCCATGCGCCGCAGTAACAATTGCTCA<br>AGCAGATTTATCGCCAGCAGCTCCGAATAGCGCCCTTCCCCTTGCCCGGCGTTAATGAT<br>TTGCCCAAACAGGTCGCTGAAATGCGGCTGGTGCGCTTCATCCGGGCGAAAGAACCCC<br>GTATTGGCAAATATTGACGGCCAGTTAAGCCATTCATGCCAGTAGGCGCGCGGACGAAA<br>GTAAACCCACTGGTGATACCATTCGCGAGCCTCCGGATGACGACCGTAGTGATGAATCT<br>CTCCTGGCGGGAACAGCAAAATATCACCCGGTCGGCAAACAAATTCTCGTCCCTGATTT<br>TTCACCACCCCCTGACCGCGAATGGTGAGATTGAGAATATAACCTTTCATTCCCAGCGGT<br>CGGTCGATAAAAAAATCGAGATAACCGTTGGCCTCAATCGGCGTTAAACCCGCCACCAG<br>ATGGGCATTAAACGAGTATCCCGGCAGCAGGGGATCATTTTGCGCTTCAGCCATactttcat<br>actcccgccattcagagaagaaaccaattgtccatattgcatcagacattgccgtcactgcgtcttttactggctcttctcgctaacca<br>aaccggtaaccccgcttattaaaagcattctgtaacaaagcgggaccaaagccatgacaaaaacgcgtaacaaaagtgtctata<br>atcacggcagaaaagtccacattgattatttgcacggcgtcacactttgctatgccatagcatttttatccataagattagcggatccta<br>cctgacgctttttatcgcaactctctactgtttctccatacccgtttttttgggctagcccacaacggtttccctctagaaataattttgtttaa<br>ctttaagaaggagatatacatATGAGCAAAGGTGAAGAACTGTTTACCGGCGTTGTGCCGATTCTG<br>GTGGAACTGGATGGCGATGTGAACGGTCACAAATTCAGCGTGCGTGGTGAAGGTGAAG<br>GCGATGCCACGATTGGCAAACTGACGCTGAAATTTATCTGCACCACCGGCAAACTGCCG</td></tr>
</table>

<table>
<tr><td>GTGCCGTGGCCGACGCTGGTGACCACCCTGACCTATGGCGTTCAGTGTTTTAGTCGCTA<br>TCCGGATCACATGAAACGTCACGATTTCTTTAAATCTGCAATGCCGGAAGGCTATGTGCA<br>GGAACGTACGATTAGCTTTAAAGATGATGGCAAATATAAAACGCGCGCCGTTGTGAAATT<br>TGAAGGCGATACCCTGGTGAACCGCATTGAACTGAAAGGCACGGATTTTAAAGAAGATG<br>GCAATATCCTGGGCCATAAACTGGAATACAACTTTAATAGCCATAATGTTTATATTACGGC<br>GGATAAACAGAAAAATGGCATCAAAGCGAATTTTACCGTTCGCCATAACGTTGAAGATGG<br>CAGTGTGCAGCTGGCAGATCATTATCAGCAGAATACCCCGATTGGTGATGGTCCGGTGC<br>TGCTGCCGGATAATCATTATCTGAGCACGCAGACCGTTCTGTCTAAAGATCCGAACGAAA<br>AAGGCACGCGGGACCACATGGTTCTGCACGAATATGTGAATGCGGCAGGTATTACGTG<br>GAGCCATCCGCAGTTCGAAAAATAAgtcgaccggctgctaacaaagcccgaaaggaagctgagttggctgctg<br>ccaccgctgagcaataactagcataaccccttggggcctctaaacgggtcttgaggggttttttg</td></tr>
</table>

<table>
<tr><td>$P_{T7}$-TriggerB</td><td>Linear DNA encoding expression of RNA trigger B under a $P_{T7}$ Promoter. Additional nucleotides (~30bp) are included in 5’ and 3’ end as extra protection against nuclease degradation.</td></tr>
<tr><td colspan="2">$P_{T7}$ Promoter-TriggerB-T7 Terminator<br><br>aacgccagcaacgcgatcccgcgaaattaatacgactcactatagggagaGGGATGCCCGTAGTTCATTCTAC<br>GGGCATGAATAACGACATACAGCAAGCGATTTACTTATACTAtagcataaccccttggggcctctaaa<br>cgggtcttgaggggttttttgtttgctgaaagccaattctga</td></tr>
</table>

<table>
<tr><td>$P_{T7}$-SwitchB-GFP</td><td>Plasmid encoding superfolder GFP expression under $P_{T7}$ promoter and toehold switch B</td></tr>
<tr><td colspan="2">$P_{T7}$ Promoter-switchB-sfGFP-T7 Terminator<br><br>taatacgactcactatagggagaGGGTATAAGTAAATCGCTTGCTGTATGTCGTTAAACAGAGGAG<br>ATAACGAATGACAGCAAGCAACCTGGCGGCAGCGCAAAAGATGAGCAAAGGTGAAGAA<br>CTGTTTACCGGCGTTGTGCCGATTCTGGTGGAACTGGATGGCGATGTGAACGGTCACAA<br>ATTCAGCGTGCGTGGTGAAGGTGAAGGCGATGCCACGATTGGCAAACTGACGCTGAAAT<br>TTATCTGCACCACCGGCAAACTGCCGGTGCCGTGGCCGACGCTGGTGACCACCCTGAC<br>CTATGGCGTTCAGTGTTTTAGTCGCTATCCGGATCACATGAAACGTCACGATTTCTTTAA<br>ATCTGCAATGCCGGAAGGCTATGTGCAGGAACGTACGATTAGCTTTAAAGATGATGGCA<br>AATATAAAACGCGCGCCGTTGTGAAATTTGAAGGCGATACCCTGGTGAACCGCATTGAA<br>CTGAAAGGCACGGATTTTAAAGAAGATGGCAATATCCTGGGCCATAAACTGGAATACAAC<br>TTTAATAGCCATAATGTTTATATTACGGCGGATAAACAGAAAAATGGCATCAAAGCGAATT<br>TTACCGTTCGCCATAACGTTGAAGATGGCAGTGTGCAGCTGGCAGATCATTATCAGCAG<br>AATACCCCGATTGGTGATGGTCCGGTGCTGCTGCCGGATAATCATTATCTGAGCACGCA<br>GACCGTTCTGTCTAAAGATCCGAACGAAAAAGGCACGCGGGACCACATGGTTCTGCACG<br>AATATGTGAATGCGGCAGGTATTACGTGGAGCCATCCGCAGTTCGAAAAATAAgtcgaccgg<br>ctgctaacaaagcccgaaaggaagctgagttggctgctgccaccgctgagcaataactagcataaccccttggggcctctaaac<br>gggtcttgaggggttttttg</td></tr>
</table>

<table>
<tr><td>$P_{T7}$-TriggerH</td><td>Linear DNA encoding expression of RNA trigger H under a $P_{T7}$ Promoter. Additional nucleotides (~30bp) are included in 5’ and 3’ end as extra protection against nuclease degradation.</td></tr>
<tr><td colspan="2">$P_{T7}$ Promoter-TriggerH-T7 Terminator</td></tr>
</table>

| |
|---|
| aacgccagcaacgcgatcccgcgaaattaatacgactcactatagggagaGGGACCGTGGACCGCATGAGGT<br>CCACGGTAAACATAACTATAACAAGCCTACAATTCATTCAAACtagcataaccccttggggcctctaaa<br>cgggtcttgaggggttttttgtttgctgaaagccaattctga |

| **$P_{T7}$-SwitchH-GFP** | Plasmid encoding superfolder GFP expression under $P_{T7}$ promoter and toehold switch H |
|---|---|
| $P_{T7}$ Promoter-switchH-sfGFP-T7 Terminator<br><br>taatacgactcactatagggagaGGGTGAATGAATTGTAGGCTTGTTATAGTTATGAACAGAGGAG<br>ACATAACATGAACAAGCCTAACCTGGCGGCAGCGCAAAAGATGAGCAAAGGTGAAGAAC<br>TGTTTACCGGCGTTGTGCCGATTCTGGTGGAACTGGATGGCGATGTGAACGGTCACAAA<br>TTCAGCGTGCGTGGTGAAGGTGAAGGCGATGCCACGATTGGCAAACTGACGCTGAAATT<br>TATCTGCACCACCGGCAAACTGCCGGTGCCGTGGCCGACGCTGGTGACCACCCTGACC<br>TATGGCGTTCAGTGTTTTAGTCGCTATCCGGATCACATGAAACGTCACGATTTCTTTAAAT<br>CTGCAATGCCGGAAGGCTATGTGCAGGAACGTACGATTAGCTTTAAAGATGATGGCAAA<br>TATAAAACGCGCGCCGTTGTGAAATTTGAAGGCGATACCCTGGTGAACCGCATTGAACT<br>GAAAGGCACGGATTTTAAAGAAGATGGCAATATCCTGGGCCATAAACTGGAATACAACTT<br>TAATAGCCATAATGTTTATATTACGGCGGATAAACAGAAAAATGGCATCAAAGCGAATTTT<br>ACCGTTCGCCATAACGTTGAAGATGGCAGTGTGCAGCTGGCAGATCATTATCAGCAGAA<br>TACCCCGATTGGTGATGGTCCGGTGCTGCTGCCGGATAATCATTATCTGAGCACGCAGA<br>CCGTTCTGTCTAAAGATCCGAACGAAAAAGGCACGCGGGACCACATGGTTCTGCACGAA<br>TATGTGAATGCGGCAGGTATTACGTGGAGCCATCCGCAGTTCGAAAAATAAgtcgaccggctg<br>ctaacaaagcccgaaaggaagctgagttggctgctgccaccgctgagcaataactagcataaccccttggggcctctaaacggg<br>tcttgaggggttttttg | |

| **$P_{T7}$-ZntR** | Plasmid encoding ZntR expression under $P_{T7}$ promoter with strong ribosomal binding site |
|---|---|
| $P_{T7}$ Promoter-Stability Hairpin-StrongRBS-ZntR-T7 Terminator<br><br>taatacgactcactatagggagaccacaacggtttccctctagaaataattttgtttaactttaagaaggagatatacatATGTAT<br>CGCATTGGTGAGCTGGCAAAAATGGCGGAAGTAACACCCGACACGATTCGTTATTACGA<br>AAAACAGCAGATGATGGAGCATGAAGTGCGTACTGAAGGTGGGTTTCGCCTATATACCG<br>AAAGCGATCTCCAGCGATTGAAATTTATCCGCCATGCCAGACAACTAGGTTTCAGTCTGG<br>AGTCGATCCGCGAGTTGCTGTCGATCCGCATCGATCCTGAACACCATACCTGTCAGGAG<br>TCAAAAGGCATTGTGCAGGAAAGATTGCAGGAAGTCGAAGCACGGATAGCCGAGTTGCA<br>GAGTATGCAGCGTTCCTTGCAACGCCTTAACGATGCCTGTTGTGGGACTGCTCATAGCA<br>GTGTTTATTGTTCGATTCTTGAAGCTCTTGAACAAGGGGCGAGTGGCGTTAAGAGTGGTT<br>GTTGATAAgtcgaccggctgctaacaaagcccgaaaggaagctgagttggctgctgccaccgctgagcaataactagcat<br>aaccccttggggcctctaaacgggtcttgaggggttttttg | |

| **$P_{zntA}$-GFP** | Plasmid encoding superfolder GFP expression under ZntR regulated $P_{zntA}$ promoter |
|---|---|
| $P_{zntA}$-Stability Hairpin-StrongRBS-sfGFP- T7 Terminator<br><br>CTGTATCTCTGATAAAACTTGACTCTGGAGTCGACTCCAGAGTGTATCCTTCGGTTAAT<br>ccacaacggtttccctctagaaataattttgtttaactttaagaaggagatatacatATGAGCAAAGGTGAAGAACTGT<br>TTACCGGCGTTGTGCCGATTCTGGTGGAACTGGATGGCGATGTGAACGGTCACAAATTC | |

AGCGTGCGTGGTGAAGGTGAAGGCGATGCCACGATTGGCAAACTGACGCTGAAATTTAT
CTGCACCACCGGCAAACTGCCGGTGCCGTGGCCGACGCTGGTGACCACCCTGACCTAT
GGCGTTCAGTGTTTTAGTCGCTATCCGGATCACATGAAACGTCACGATTTCTTTAAATCT
GCAATGCCGGAAGGCTATGTGCAGGAACGTACGATTAGCTTTAAAGATGATGGCAAATA
TAAAACGCGCGCCGTTGTGAAATTTGAAGGCGATACCCTGGTGAACCGCATTGAACTGA
AAGGCACGGATTTTAAAGAAGATGGCAATATCCTGGGCCATAAACTGGAATACAACTTTA
ATAGCCATAATGTTTATATTACGGCGGATAAACAGAAAAATGGCATCAAAGCGAATTTTAC
CGTTCGCCATAACGTTGAAGATGGCAGTGTGCAGCTGGCAGATCATTATCAGCAGAATA
CCCCGATTGGTGATGGTCCGGTGCTGCTGCCGGATAATCATTATCTGAGCACGCAGACC
GTTCTGTCTAAAGATCCGAACGAAAAAGGCACGCGGACCACATGGTTCTGCACGAATA
TGTGAATGCGGCAGGTATTACGTGGAGCCATCCGCAGTTCGAAAAATAAgtcgaccggctgcta
acaaagcccgaaaggaagctgagttggctgctgccaccgctgagcaataactagcataaccccttggggcctctaaacgggtctt
gaggggttttttg

| **$P_{T7}$-EutR** | Plasmid encoding EutR expression under $P_{T7}$ promoter with strong ribosomal binding site. |
|---|---|

$P_{T7}$ Promoter-Stability Hairpin-StrongRBS-EutS-T7 Terminator

taatacgactcactatagggagaccacaacggtttccctctagaaataattttgtttaactttaagaaggagatatacatATGAAA
AAGACCCGTACAGCCAATTTGCACCATCTTTATCATGAACCCTTACCCGAAAACCTGAAG
CTCACGCCGAAGGTCGAAGTGGATAATGTTCATCAACGACAGACAACGGATGTCTATGA
ACATGCTTTAACGATTACCGCCTGGCAGCAGATTTACGATCAGCTGCATCCGGGCAAGT
TTCATGGTGAATTTACGGAAATTCTACTCGATGATATTCAGGTTTTTCGTGAATACACCGG
TCTGGCGCTGCGTCAGTCGTGCCTGGTCTGGCCGAACTCGTTCTGGTTTGGCATTCCGG
CGACGCGCGGTGAGCAGGGATTTATCGGTTCGCAATGTCTGGGAAGCGCGGAAATCGC
CACCCGCCCTGGTGGCACTGAATTTGAACTGAGCACGCCGGATGATTACACGATCCTGG
GCGTGGTGCTTTCTGAAGATGTCATCACCCGGCAGGCTAACTTTTTGCATAACCCGGAT
CGGGTATTACATATGTTGCGTAACCAGTCGGCGCTGGAAGTGAAAGAGCAGCATAAAGC
CGCGCTGTGGGGCTTTGTCCAACAGGCGCTGGCGACGTTTTGCGAGAATCCGGAAAAT
CTCCATCAGCCAGCAGTGCGAAAAGTGCTGGGGGATAATTTGCTAATGGCGATGGGGG
CCATGCTGGAAGAAGCGCAACCAATGGTGACGGCGGAAAGCATCAGTCATCAGAGTTAC
CGTCGATTGCTTTCCCGCGCCCGTGAATATGTGCTGGAAAACATGTCCGAACCGGTGAC
GGTGCTGGATTTGTGTAATCAACTGCATGTCAGCCGCCGCACGCTACAAAACGCGTTTC
ACGCTATTTTAGGCATTGGCCCGAACGCGTGGCTGAAACGCATTCGCCTGAACGCCGTA
CGCCGCGAACTGATAAGTCCGTGGTCGCAAAGTATGACGGTAAAAGACGCCGCCATGC
AGTGGGGATTCTGGCATCTGGGGCAATTTGCCACGGATTACCAGCAGCTGTTTTCCGAG
AAGCCGTCACTGACGCTGCATCAGCGGATGCGGGAGTGGGGGTGAgtcgaccggctgctaaca
aagcccgaaaggaagctgagttggctgctgccaccgctgagcaataactagcataaccccttggggcctctaaacgggtcttga
ggggttttttg

| **$P_{eutS}$-GFP** | Plasmid encoding superfolder GFP expression under EutR regulated $P_{eutS}$ promoter |
|---|---|

$P_{eutS}$-Stability Hairpin-StrongRBS-sfGFP- T7 Terminator

AACAGAGCGAAGTGGTGGCTTCAGCGTATGGCGATCAGGATCTGAGCTTTGGTCCGGA
ATACATCATTCCAAAACCGTTTGATCCGCGCTTGATCGTTAAGATCGCTCCTGCGGTCGC
TAAAGCCGCGATGGAGTCGGGCGTGGCGACTCGTCCGATTGCTGATTTCGACGTCTACA
TCGACAAGCTGACTGAGTTCGTTTACAAAACCAACCTGTTTATGAAGCCGATTTTCTCCC

AGGCTCGCAAAGCGCCGAAGCGCGTTGTTCTGCCGGAAGGGGAAGAGGCGCGCGTTC
TGCATGCCACTCAGGAACTGGTAACGCTGGGACTGGCGAAACCGATCCTTATCGGTCGT
CCGAACGTGATCGAAATGCGCATTCAGAAACTGGGCTTGCAGATCAAAGCGAGCGTTGA
TTTTGAGATCGTCAATAACGAATCCGATCCGCGCTTTAAAGAGTACTGGACCGAATACTT
CCAGATCATGAAGCGTCGCGGCGTCACTCAGGAACAGGCGCAGCGGGCGCTGATCAGT
AACCCGACAGTGATCGGCGCGATCATGGTTCAGCGTGGGGAAGCCGATGCAATGATTT
GCGGTACGGTGGGTGATTATCATGAACATTTTAGCGTGGTGAAAAATGTCTTTGGTTATC
GCGATGGCGTTCACACCGCAGGTGCCATGAACGCGCTGCTGCTGCCGAGTGGTAACAC
CTTTATTGCCGATACATATGTTAATGATGAACCGGATGCAGAAGAGCTGGCGGAGATCA
CCTTGATGGCGGCAGAAACTGTCCGTCGTTTTGGTATTGAGCCGCGCGTTGCTTTGTTG
TCGCACTCCAACTTTGGTTCTTCTGACTGCCCGTCGTCGAGCAAAATGCGTCAGGCGCT
GGAACTGGTCAGGGAACGTGCACCAGAACTGATGATTGATGGTGAAATGCACGGCGAT
GCAGCGCTGGTGGAAGCGATTCGCAACGACCGTATGCCGGACAGCTCTTTGAAAGGTT
CCGCCAATATTCTGGTGATGCCGAACATGGAAGCTGCCCGCATTAGTTACAACTTACTG
CGTGTTTCCAGCTCGGAAGGTGTGACTGTCGGCCCGGTGCTGATGGGTGTGGCGAAAC
CGGTTCACGTGTTAACGCCGATCGCATCGGTGCGTCGTATCGTCAACATGGTGGCGCTG
GCCGTGGTAGAAGCGCAAACCCAACCGCTGTAATTTTTTTTAACTCTCACGCTTATCCTG
AATATTCAGGGTAAGCAGTTTAGCTGCAATATATTAGTAAAGCTTATTACTGAGTTTGCGA
ATAATAAAAAAAAGCAGTCTATATAATATCTCGATATTATTTTATTTATATTCATGCGTTGC
ATATGAAAGTTTATGCACCACAGCGAATATCTCTCCATTCTTAGTGATCTACCTCACCTTT
TAAACGCGCTTGCCGAATTTTGTTATTTACTCTGACGAAAAATTGTCACGATACACGAAA
GTTTTTCACAGGCGGCGACTCccacaacggtttccctctagaaataattttgtttaactttaagaaggagatatacatA
TGAGCAAAGGTGAAGAACTGTTTACCGGCGTTGTGCCGATTCTGGTGGAACTGGATGGC
GATGTGAACGGTCACAAATTCAGCGTGCGTGGTGAAGGTGAAGGCGATGCCACGATTG
GCAAACTGACGCTGAAATTTATCTGCACCACCGGCAAACTGCCGGTGCCGTGGCCGAC
GCTGGTGACCACCCTGACCTATGGCGTTCAGTGTTTTAGTCGCTATCCGGATCACATGA
AACGTCACGATTTCTTTAAATCTGCAATGCCGGAAGGCTATGTGCAGGAACGTACGATTA
GCTTTAAAGATGATGGCAAATATAAAACGCGCGCCGTTGTGAAATTTGAAGGCGATACCC
TGGTGAACCGCATTGAACTGAAAGGCACGGATTTTAAAGAAGATGGCAATATCCTGGGC
CATAAACTGGAATACAACTTTAATAGCCATAATGTTTATATTACGGCGGATAAACAGAAAA
ATGGCATCAAAGCGAATTTTACCGTTCGCCATAACGTTGAAGATGGCAGTGTGCAGCTG
GCAGATCATTATCAGCAGAATACCCCGATTGGTGATGGTCCGGTGCTGCTGCCGGATAA
TCATTATCTGAGCACGCAGACCGTTCTGTCTAAAGATCCGAACGAAAAAGGCACGCGGG
ACCACATGGTTCTGCACGAATATGTGAATGCGGCAGGTATTACGTGGAGCCATCCGCAG
TTCGAAAAATAAgtcgaccggctgctaacaaagcccgaaaggaagctgagttggctgctgccaccgctgagcaataact
agcataaccccttggggcctctaaacgggtcttgaggggttttttg

| **$P_{T7}$-*B. theta* Trigger** | Linear DNA encoding expression of RNA *B. theta* trigger under a $P_{T7}$ Promoter. Additional nucleotides (~30bp) are included in 5' and 3' end as extra protection against nuclease degradation. |
|---|---|

$P_{T7}$ Promoter-B. theta Trigger

ggaaaaacgccagcaacgcgatcccgcgaaattaatacgactcactataggCCGACTTCGGAACGCTTATAGA
AAGGAGCAACACCACACAAAGCCGGTCATACAGTAATTCAGCTACCGCATACGTTTCAaa
aaaaaacgccgcctttcggcggcgtttg

<table>
<tr><td>$P_{T7}$-B. theta Switch-GFP</td><td>Plasmid encoding superfolder GFP expression under $P_{T7}$ promoter and B. theta toehold switch</td></tr>
<tr><td colspan="2">T7 Promoter-B. theta Switch-sfGFP-T7 Terminator<br><br>taatacgactcactatagggagaGTTACTGTATGACCGGCTTTGTGTGGTGTTGCTCCTTGGACTTT<br>AGAACAGAGGAGATAAAGATGAAGGAGCAACACAACCTGGCGGCAGCGCAAAAGATGA<br>GCAAAGGTGAAGAACTGTTTACCGGCGTTGTGCCGATTCTGGTGGAACTGGATGGCGAT<br>GTGAACGGTCACAAATTCAGCGTGCGTGGTGAAGGTGAAGGCGATGCCACGATTGGCA<br>AACTGACGCTGAAATTTATCTGCACCACCGGCAAACTGCCGGTGCCGTGGCCGACGCT<br>GGTGACCACCCTGACCTATGGCGTTCAGTGTTTTAGTCGCTATCCGGATCACATGAAAC<br>GTCACGATTTCTTTAAATCTGCAATGCCGGAAGGCTATGTGCAGGAACGTACGATTAGCT<br>TTAAAGATGATGGCAAATATAAAACGCGCGCCGTTGTGAAATTTGAAGGCGATACCCTG<br>GTGAACCGCATTGAACTGAAAGGCACGGATTTTAAAGAAGATGGCAATATCCTGGGCCA<br>TAAACTGGAATACAACTTTAATAGCCATAATGTTTATATTACGGCGGATAAACAGAAAAAT<br>GGCATCAAAGCGAATTTTACCGTTCGCCATAACGTTGAAGATGGCAGTGTGCAGCTGGC<br>AGATCATTATCAGCAGAATACCCCGATTGGTGATGGTCCGGTGCTGCTGCCGGATAATC<br>ATTATCTGAGCACGCAGACCGTTCTGTCTAAAGATCCGAACGAAAAAGGCACGCGGGAC<br>CACATGGTTCTGCACGAATATGTGAATGCGGCAGGTATTACGTGGAGCCATCCGCAGTT<br>CGAAAAATAAgtcgaccggctgctaacaaagcccgaaaggaagctgagttggctgctgccaccgctgagcaataactag<br>cataaccccttggggcctctaaacgggtcttgaggggttttttg</td></tr>
</table>

<table>
<tr><td>$P_{T7}$-Stx1 Trigger</td><td>Linear DNA encoding expression of RNA Stx1 trigger under a $P_{T7}$ Promoter. Additional nucleotides (~30bp) are included in 5’ and 3’ end as extra protection against nuclease degradation.</td></tr>
<tr><td colspan="2">$P_{T7}$ Promoter-Stx1 Trigger<br><br>ggaaaaacgccagcaacgcgatcccgcgaaattaatacgactcactataggATAAATCGCCATTCGTTGACTAC<br>TTCTTATCTGGATTTAATGTCGCATAGTGGAACCTCACTGACGCAGTCTGTGGCAAGAGC<br>GATGTTACGGTTTaaaaaaaacgccgcctttcggcggcgtttg</td></tr>
</table>

<table>
<tr><td>$P_{T7}$-Stx1 Switch-GFP</td><td>Plasmid encoding superfolder GFP expression under $P_{T7}$ promoter and Stx1 toehold switch</td></tr>
<tr><td colspan="2">$P_{T7}$ Promoter-Stx1 Switch-sfGFP-T7 Terminator<br><br>taatacgactcactatagggagaGGGCGTCAGTGAGGTTCCACTATGCGACATTAAATCCAGGGAC<br>TTTAGAACAGAGGAGATAAAGATGCTGGATTTAATTAACCTGGCGGCAGCGCAAAAGAT<br>GAGCAAAGGTGAAGAACTGTTTACCGGCGTTGTGCCGATTCTGGTGGAACTGGATGGC<br>GATGTGAACGGTCACAAATTCAGCGTGCGTGGTGAAGGTGAAGGCGATGCCACGATTG<br>GCAAACTGACGCTGAAATTTATCTGCACCACCGGCAAACTGCCGGTGCCGTGGCCGAC<br>GCTGGTGACCACCCTGACCTATGGCGTTCAGTGTTTTAGTCGCTATCCGGATCACATGA<br>AACGTCACGATTTCTTTAAATCTGCAATGCCGGAAGGCTATGTGCAGGAACGTACGATTA<br>GCTTTAAAGATGATGGCAAATATAAAACGCGCGCCGTTGTGAAATTTGAAGGCGATACCC<br>TGGTGAACCGCATTGAACTGAAAGGCACGGATTTTAAAGAAGATGGCAATATCCTGGGC<br>CATAAACTGGAATACAACTTTAATAGCCATAATGTTTATATTACGGCGGATAAACAGAAAA<br>ATGGCATCAAAGCGAATTTTACCGTTCGCCATAACGTTGAAGATGGCAGTGTGCAGCTG<br>GCAGATCATTATCAGCAGAATACCCCGATTGGTGATGGTCCGGTGCTGCTGCCGGATAA<br>TCATTATCTGAGCACGCAGACCGTTCTGTCTAAAGATCCGAACGAAAAAGGCACGCGGG</td></tr>
</table>

<table>
<tr><td>ACCACATGGTTCTGCACGAATATGTGAATGCGGCAGGTATTACGTGGAGCCATCCGCAG<br>TTCGAAAAATAAgtcgaccggctgctaacaaagcccgaaaggaagctgagttggctgctgccaccgctgagcaataact<br>agcataaccccttggggcctctaaacgggtcttgaggggttttttg</td></tr>
</table>

<table>
<tr><th>$P_{T7}$-Stx2 Trigger</th><td>Linear DNA encoding expression of RNA Stx2 trigger under a $P_{T7}$ Promoter. Additional nucleotides (~30bp) are included in 5’ and 3’ end as extra protection against nuclease degradation.</td></tr>
<tr><td colspan="2">$P_{T7}$ Promoter-Stx2 Trigger<br><br>ggaaaaacgccagcaacgcgatcccgcgaaattaatacgactcactataggGTATCCTATTCCCGGGAGTTTAC<br>GATAGACTTTTCGACCCAACAAAGTTATGTCTCTTCGTTAAATAGTATACGGACAGAGATA<br>TCGACCCCTCTTGAACATATATCaaaaaaaacgccgcctttcggcggcgtttg</td></tr>
</table>

<table>
<tr><th>$P_{T7}$-Stx2 Switch-GFP</th><td>Plasmid encoding superfolder GFP expression under $P_{T7}$ promoter and Stx2 toehold switch</td></tr>
<tr><td colspan="2">$P_{T7}$ Promoter-Stx2 Switch-sfGFP-T7 Terminator<br><br>taatacgactcactatagggagaGGGATACTATTTAACGAAGAGACATAACTTTGTTGGGTCGGACT<br>TTAGAACAGAGGAGATAAAGATGGACCCAACAAAGATGAGCAAAGGTGAAGAACTGTTT<br>ACCGGCGTTGTGCCGATTCTGGTGGAACTGGATGGCGATGTGAACGGTCACAAATTCAG<br>CGTGCGTGGTGAAGGTGAAGGCGATGCCACGATTGGCAAACTGACGCTGAAATTTATCT<br>GCACCACCGGCAAACTGCCGGTGCCGTGGCCGACGCTGGTGACCACCCTGACCTATGG<br>CGTTCAGTGTTTTAGTCGCTATCCGGATCACATGAAACGTCACGATTTCTTTAAATCTGCA<br>ATGCCGGAAGGCTATGTGCAGGAACGTACGATTAGCTTTAAAGATGATGGCAAATATAAA<br>ACGCGCGCCGTTGTGAAATTTGAAGGCGATACCCTGGTGAACCGCATTGAACTGAAAGG<br>CACGGATTTTAAAGAAGATGGCAATATCCTGGGCCATAAACTGGAATACAACTTTAATAG<br>CCATAATGTTTATATTACGGCGGATAAACAGAAAAATGGCATCAAAGCGAATTTTACCGTT<br>CGCCATAACGTTGAAGATGGCAGTGTGCAGCTGGCAGATCATTATCAGCAGAATACCCC<br>GATTGGTGATGGTCCGGTGCTGCTGCCGGATAATCATTATCTGAGCACGCAGACCGTTC<br>TGTCTAAAGATCCGAACGAAAAAGGCACGCGGGACCACATGGTTCTGCACGAATATGTG<br>AATGCGGCAGGTATTACGTGGAGCCATCCGCAGTTCGAAAAATAAgtcgaccggctgctaacaaa<br>gcccgaaaggaagctgagttggctgctgccaccgctgagcaataactagcataaccccttggggcctctaaacgggtcttgagg<br>ggttttttg</td></tr>
</table>

<table>
<tr><th>$P_{T7}$-LacZ</th><td>Plasmid encoding β-galactosidase (LacZ) expression under $P_{T7}$ promoter</td></tr>
<tr><td colspan="2">$P_{T7}$ Promoter- Stability Hairpin-StrongRBS-LacZ-T7 Terminator<br><br>taatacgactcactatagggagaccacaacggtttccctctagaaataattttgtttaactttaagaaggagatatacatatgACCA<br>TGATTACGGATTCACTGGCCGTCGTTTTACAACGTCGTGACTGGGAAAACCCTGGCGTT<br>ACCCAACTTAATCGCCTTGCAGCACATCCCCCTTTCGCCAGCTGGCGTAATAGCGAAGA<br>GGCCCGCACCGATCGCCCTTCCCAACAGTTGCGCAGCCTGAATGGCGAATGGCGCTTT<br>GCCTGGTTTCCGGCACCAGAAGCGGTGCCGGAAAGCTGGCTGGAGTGCGATCTTCCTG<br>AGGCCGATACTGTCGTCGTCCCCTCAAACTGGCAGATGCACGGTTACGATGCGCCCATC<br>TACACCAACGTGACCTATCCCATTACGGTCAATCCGCCGTTTGTTCCCACGGAGAATCC<br>GACGGGTTGTTACTCGCTCACATTTAATGTTGATGAAAGCTGGCTACAGGAAGGCCAGA<br>CGCGAATTATTTTTGATGGCGTTAACTCGGCGTTTCATCTGTGGTGCAACGGGCGCTGG</td></tr>
</table>

GTCGGTTACGGCCAGGACAGTCGTTTGCCGTCTGAATTTGACCTGAGCGCATTTTTACG
CGCCGGAGAAAACCGCCTCGCGGTGATGGTGCTGCGCTGGAGTGACGGCAGTTATCTG
GAAGATCAGGATATGTGGCGGATGAGCGGCATTTTCCGTGACGTCTCGTTGCTGCATAA
ACCGACTACACAAATCAGCGATTTCCATGTTGCCACTCGCTTTAATGATGATTTCAGCCG
CGCTGTACTGGAGGCTGAAGTTCAGATGTGCGGCGAGTTGCGTGACTACCTACGGGTA
ACAGTTTCTTTATGGCAGGGTGAAACGCAGGTCGCCAGCGGCACCGCGCCTTTCGGCG
GTGAAATTATCGATGAGCGTGGTGGTTATGCCGATCGCGTCACACTACGTCTGAACGTC
GAAAACCCGAAACTGTGGAGCGCCGAAATCCCGAATCTCTATCGTGCGGTGGTTGAACT
GCACACCGCCGACGGCACGCTGATTGAAGCAGAAGCCTGCGATGTCGGTTTCCGCGAG
GTGCGGATTGAAAATGGTCTGCTGCTGCTGAACGGCAAGCCGTTGCTGATTCGAGGCGT
TAACCGTCACGAGCATCATCCTCTGCATGGTCAGGTCATGGATGAGCAGACGATGGTGC
AGGATATCCTGCTGATGAAGCAGAACAACTTTAACGCCGTGCGCTGTTCGCATTATCCG
AACCATCCGCTGTGGTACACGCTGTGCGACCGCTACGGCCTGTATGTGGTGGATGAAG
CCAATATTGAAACCCACGGCATGGTGCCAATGAATCGTCTGACCGATGATCCGCGCTGG
CTACCGGCGATGAGCGAACGCGTAACGCGAATGGTGCAGCGCGATCGTAATCACCCGA
GTGTGATCATCTGGTCGCTGGGGAATGAATCAGGCCACGGCGCTAATCACGACGCGCT
GTATCGCTGGATCAAATCTGTCGATCCTTCCCGCCCGGTGCAGTATGAAGGCGGCGGA
GCCGACACCACGGCCACCGATATTATTTGCCCGATGTACGCGCGCGTGGATGAAGACC
AGCCCTTCCCGGCTGTGCCGAAATGGTCCATCAAAAAATGGCTTTCGCTACCTGGAGAG
ACGCGCCCGCTGATCCTTTGCGAATACGCCCACGCGATGGGTAACAGTCTTGGCGGTTT
CGCTAAATACTGGCAGGCGTTTCGTCAGTATCCCCGTTTACAGGGCGGCTTCGTCTGGG
ACTGGGTGGATCAGTCGCTGATTAAATATGATGAAAACGGCAACCCGTGGTCGGCTTAC
GGCGGTGATTTTGGCGATACGCCGAACGATCGCCAGTTCTGTATGAACGGTCTGGTCTT
TGCCGACCGCACGCCGCATCCAGCGCTGACGGAAGCAAAACACCAGCAGCAGTTTTTC
CAGTTCCGTTTATCCGGGCAAACCATCGAAGTGACCAGCGAATACCTGTTCCGTCATAG
CGATAACGAGCTCCTGCACTGGATGGTGGCGCTGGATGGTAAGCCGCTGGCAAGCGGT
GAAGTGCCTCTGGATGTCGCTCCACAAGGTAAACAGTTGATTGAACTGCCTGAACTACC
GCAGCCGGAGAGCGCCGGGCAACTCTGGCTCACAGTACGCGTAGTGCAACCGAACGC
GACCGCATGGTCAGAAGCCGGACACATCAGCGCCTGGCAGCAGTGGCGTCTGGCTGAA
AACCTCAGCGTGACACTCCCCGCCGCGTCCCACGCCATCCCGCATCTGACCACCAGCG
AAATGGATTTTTGCATCGAGCTGGGTAATAAGCGTTGGCAATTTAACCGCCAGTCAGGCT
TTCTTTCACAGATGTGGATTGGCGATAAAAAACAACTGCTGACGCCGCTGCGCGATCAG
TTCACCCGTGCACCGCTGGATAACGACATTGGCGTAAGTGAAGCGACCCGCATTGACCC
TAACGCCTGGGTCGAACGCTGGAAGGCGGCGGGCCATTACCAGGCCGAAGCAGCGTT
GTTGCAGTGCACGGCAGATACACTTGCTGATGCGGTGCTGATTACGACCGCTCACGCGT
GGCAGCATCAGGGGAAAACCTTATTTATCAGCCGGAAAACCTACCGGATTGATGGTAGT
GGTCAAATGGCGATTACCGTTGATGTTGAAGTGGCGAGCGATACACCGCATCCGGCGC
GGATTGGCCTGAACTGCCAGCTGGCGCAGGTAGCAGAGCGGGTAAACTGGCTCGGATT
AGGGCCGCAAGAAAACTATCCCGACCGCCTTACTGCCGCCTGTTTTGACCGCTGGGATC
TGCCATTGTCAGACATGTATACCCCGTACGTCTTCCCGAGCGAAAACGGTCTGCGCTGC
GGGACGCGCGAATTGAATTATGGCCCACACCAGTGGCGCGGCGACTTCCAGTTCAACA
TCAGCCGCTACAGTCAACAGCAACTGATGGAAACCAGCCATCGCCATCTGCTGCACGCG
GAAGAAGGCACATGGCTGAATATCGACGGTTTCCATATGGGGATTGGTGGCGACGACTC
CTGGAGCCCGTCAGTATCGGCGGAATTCCAGCTGAGCGCCGGTCGCTACCATTACCAG
TTGGTCTGGTGTCAAAAAtaagtcgaccggctgctaacaaagcccgaaaggaagctgagttggctgctgccaccgct
gagcaataactagcataaccccttggggcctctaaacgggtcttgaggggttttttg

| **$\chi$DNA** | DNA oligo containing 6 $\chi$ sites to stall endonuclease degradation on linear DNA templates. |
|---|---|
| FW: TCACTTCACTGCTGGTGGCCACTGCTGGTGGCCACTGCTGGTGGCCACTGCTGGTGGCCACTGCTGGTGGCCACTGCTGGTGGCCA | |
| RV: TGGCCACCAGCAGTGGCCACCAGCAGTGGCCACCAGCAGTGGCCACCAGCAGTGGCCACCAGCAGTGGCCACCAGCAGTGAAGTGA | |

| **$P_{T7}$-Null** | Linear DNA encoding expression of random RNA sequences under a $P_{T7}$ Promoter. Additional nucleotides (~30bp) are included in 5' and 3' end as extra protection against nuclease degradation. |
|---|---|
| $P_{T7}$ Promoter-Null<br><br>ggaaaaacgccagcaacgcgatcccgcgaaattaatacgactcactataggGTCGACCGGCTGCTAACAAAGCCCGAAAGGAAGCTGAGTTGGCTGCTGCCACCGCTGAGCAATAACaaaaaaaacgccgcctttcggcggcgtttg | |

| **$P_{T7}$-*E. coli* Trigger** | Linear DNA encoding expression of RNA *E. coli* trigger under a $P_{T7}$ Promoter. Additional nucleotides (~30bp) are included in 5' and 3' end as extra protection against nuclease degradation. |
|---|---|
| $P_{T7}$ Promoter-*E. coli* Trigger<br><br>ggaaaaacgccagcaacgcgatcccgcgaaattaatacgactcactataggAAGACGGATATCTATTTCGTTTCCACGTTTGAACCGaaaaaaaacgccgcctttcggcggcgtttg | |

**Table S2.** Description of lysate, plasmid concentrations, and reaction additives present in protocell sensors or CFE reactions in each figure.

| Figures | Detecting | Lysate Preparation | Plasmids | Reaction Additives |
|---|---|---|---|---|
| 2, S2 | | Basal level T7 RNAP | 1 nM $P_{T7}$-GFP | |
| 3 | IPTG | Basal level T7 RNAP | 5 nM $P_{T7}$-LacI<br>2.5 nM $P_{T7lacO}$-GFP | |
| 3 | Arabinose | Basal level T7 RNAP | 10 nM AraC-$P_{BAD}$-GFP | |
| 4C, D, S3 | RNA B | Enriched in T7 RNAP | 2.5 nM $P_{T7}$-switchB-GFP | |
| 4C, D, S3 | RNA H | Enriched in T7 RNAP | 2.5 nM $P_{T7}$-switchH-GFP | |
| 4E, F, S4, S5 | Linear DNA B Trigger | Enriched in T7 RNAP | 2.5 nM $P_{T7}$-switchB-GFP | 2 $\mu$M $\chi$DNA |
| 4E, F, S4, S5 | Linear DNA H Trigger | Enriched in T7 RNAP | 2.5 nM $P_{T7}$-switchH-GFP | 2 $\mu$M $\chi$DNA |
| 5B, S6 | Zinc | Basal level T7 RNAP | 0.25 nM $P_{T7}$-ZntR<br>2.5 nM $P_{ZntA}$-GFP | |
| 5B, C, S6 | Vitamin $B_{12}$ | Basal level T7 RNAP | 2.5 nM $P_{T7}$-EutR<br>10 nM $P_{EutS}$-GFP | |
| 5B, C | RNA Stx1 Trigger | Enriched in T7 RNAP | 4 nM $P_{T7}$-Stx1 switch-GFP | |
| 5B, C, S7 | Linear DNA Stx2 Trigger | Enriched in T7 RNAP | 5 nM $P_{T7}$-Stx2 switch-GFP | 2 $\mu$M $\chi$DNA |
| 5C | Zinc | Basal level T7 RNAP | 0.10 nM $P_{T7}$-ZntR<br>2.5 nM $P_{ZntA}$-GFP | |
| 6A | Linear DNA *B. theta* Trigger | Enriched in T7 RNAP | 1.5 nM $P_{T7}$-*B. theta* switch-LacZ | 2 $\mu$M $\chi$DNA |
| 6A | Linear DNA Stx1 Trigger | Enriched in T7 RNAP | 1 nM $P_{T7}$-Stx1-LacZ | 2 $\mu$M GamS |
| 6A | Linear DNA Stx2 Trigger | Enriched in T7 RNAP | 1 nM $P_{T7}$-Stx2-LacZ | 2 $\mu$M $\chi$DNA |
| 6B | Linear DNA *B. theta* Trigger | Enriched in T7 RNAP | 3 nM $P_{T7}$-*B. theta* switch-LacZ | 2 $\mu$M $\chi$DNA |
| 6B | Linear DNA Stx1 Trigger | Enriched in T7 RNAP | 3 nM $P_{T7}$-Stx1-LacZ | 2 $\mu$M GamS |
| 6B | Linear DNA Stx2 Trigger | Enriched in T7 RNAP | 3 nM $P_{T7}$-Stx2-LacZ | 2 $\mu$M $\chi$DNA |
| S7 | Linear DNA *B. theta* Trigger | Enriched in T7 RNAP | 5 nM $P_{T7}$-*B. theta* switch-GFP | 2 $\mu$M $\chi$DNA |

| | | | | |
|---|---|---|---|---|
| S7 | Linear DNA<br>Stx1 Trigger | Enriched in<br>T7 RNAP | 4 nM $P_{T7}$-*B. theta* switch-GFP | 2 $\mu$M GamS |
| S8 | | Basal level<br>T7 RNAP | 2 nM $P_{T7}$-LacZ | |
| S10 | Plasmid<br>*B. theta* Trigger | Enriched in<br>T7 RNAP | 5 nM $P_{T7}$-*B. theta* trigger<br>2.5 nM $P_{T7}$-*B. theta* switch-GFP | |
| S10 | Plasmid Stx1<br>Trigger | Enriched in<br>T7 RNAP | 5 nM $P_{T7}$-Stx1 trigger<br>2.5 nM $P_{T7}$-Stx1 switch-GFP | |
| S10 | Plasmid Stx2<br>Trigger | Enriched in<br>T7 RNAP | 5 nM $P_{T7}$-Stx2 trigger<br>2.5 nM $P_{T7}$-Stx2 switch-GFP | |

**Table S3:** Primers used for trigger DNA amplification. Lowercase, unlabeled sequences are protective regions to decrease endonuclease degradation. Highlighted sequences indicate the T7 promoter, and uppercase sequences are primer annealing regions. Target-specific primers were designed bind to 30-35 base pairs before and after the actual trigger sequence to prevent unintended primer activation of switches. The specificity of the developed switches was also validated on the genomic DNA of STEC O157: H7, *B. thetaiotaomicron,* and common laboratory *E. coli* strains DH10β and BL21 Star (DE3) (**Figure S10**).

| Amplifying | Primer Sequence |
|---|---|
| B or H | Fwd:<br>aacgccagcaacgcgatcccgcgaaatTAATACGACTCACTATAGGGAGA<br><br>Rev:<br>TaatcagaattggctttcagcaaaAAACCCCTCAAGACCCGTT<br><br>Template: T7-triggerB or T7-triggerH |
| *B. theta* | Fwd:<br>ggaaaaacgccagcaacgcgatcccgcgaaattaatacgactcactataggCCGACTTCGGAACGCTTATAGA<br><br>Rev:<br>caaacgccgccgaaaggcggcgttttttttTGAAACGTATGCGGTAGCTGAA<br><br>Genomic Target: hypothetical protein SAMN029103 22_01913<br>ATGCATGCATACATTATCCAACAACTAACAAGAATTATATTGTTTATCACTATCGGTTTGCCTATAGGACTAAAAAGTTTTGCCCAAGAAACAAAACGTTTCTATATGGAACTGGACACTCCCCGCAATGGAGCCAAAGCAGGACAAGAGCTTGAATTAAAATACATCAGCACAGCCGATTTCGATTCTGTATCTCCACCCGACTTCGGAACGCTTATAGAAACAGTTGAAGGAGCAACACCACACAAAGCCGGTCATACAGTAAAAAACGGCATATTGACAGATATCTACGAGCAGGGATTCAGCTACCGCATACGTTTCAAGAAGCCAGGAAACACCAAACTACCTCTGGCATCCATCAAGGCAAACGGAAAGGAATACGAAACACCTCTGACCAGTGTATGGGTACATCCGGTCGATACCAATATCGACAGTGTAAAATGCAGCATTCAGCTGGAGGATTCTTATCGCAAAGGAGTTTTCACTGCCATCGGGATCTGTCTCTTAATCGCCTGGTTATTGATCCGCTTATCGTTTCAGAAACAAAAAAATAAAGAGACAGGATAA |
| Stx1 | Fwd:<br>ggaaaaacgccagcaacgcgatcccgcgaaattaatacgactcactataggATAAATCGCCATTCGTTGACTACT<br><br>Rev:<br>caaacgccgccgaaaggcggcgttttttttAAACCGTAACATCGCTCTTGCCA<br><br>Genomic Target: Shiga Toxin 1<br>ATGAAAATAATTATTTTTAGAGTGCTAACTTTTTTCTTTGTTATCTTTTCAGTTAATGTGGTTGCGAAGGAATTTACCTTAGACTTCTCGACTGCAAAGACGTATGTAGATTCGCTGAATGTCATTCGCTCTGCAATAGGTACTCCATTACAGACTATTTCATCAGGAGGTACGTCTTTACTGATGATTGATAGTGGCACAGGG |

| | |
|---|---|
| | GATAATTTGTTTGCAGTTGATGTCAGAGGGATAGATCCAGAGGAAGGGCG<br>GTTTAATAATCTACGGCTTATTGTTGAACGAAATAATTTATATGTGACAGG<br>ATTTGTTAACAGGACAAATAATGTTTTTTATCGCTTTGCTGATTTTTCACAT<br>GTTACCTTTCCAGGTACAACAGCGGTTACATTGTCTGGTGACAGTAGCTA<br>TACCACGTTACAGCGTGTTGCAGGGATCAGTCGTACGGGGATGCAGATA<br>AATCGCCATTCGTTGACTACTTCTTATCTGGATTTAATGTCGCATAGTGGA<br>ACCTCACTGACGCAGTCTGTGGCAAGAGCGATGTTACGGTTTGTTACTGT<br>GACAGCTGAAGCTTTACGTTTTCGGCAAATACAGAGGGGATTTCGTACAA<br>CACTGGATGATCTCAGTGGGCGTTCTTATGTAATGACTGCTGAAGATGTT<br>GATCTTACATTGAACTGGGGAAGGTTGAGTAGTGTCCTGCCTGATTATCA<br>TGGACAAGACTCTGTTCGTGTAGGAAGAATTTCTTTTGGAAGCATTAATGC<br>AATTCTGGGAAGCGTGGCATTAATACTGAATTGTCATCATCATGCATCGC<br>GAGTTGCCAGAATGGCATCTGATGAGTTTCCTTCTATGTGTCCGGCAGAT<br>GGAAGAGTCCGTGGGATTACGCACAATAAAATATTGTGGGATTCATCCAC<br>TCTGGGGGCAATTCTGATGCGCAGAACTATTAGCAGTTG |
| Stx2 | Fwd:<br>ggaaaaacgccagcaacgcgatcccgcgaaattaatacgactcactatagGGTATCCTATTCCC<br>GGGAGTTTACGATAGACTTTTC<br><br>Rev:<br>caaacgccgccgaaaggcggcgttttttttGATATATGTTCAAGAGGGGTCGATATCTCT<br>GTCCG<br><br>Genomic Target: Shiga Toxin 2<br>ATGAAGTGTATATTATTTAAATGGGTACTGTGCCTGTTACTGGGTTTTTCTT<br>CGGTATCCTATTCCCGGGAGTTTACGATAGACTTTTCGACCCAACAAAGT<br>TATGTCTCTTCGTTAAATAGTATACGGACAGAGATATCGACCCCTCTTGAA<br>CATATATCTCAGGGGACCACATCGGTGTCTGTTATTAACCACACCCCACC<br>GGGCAGTTATTTTGCTGTGGATATACGAGGGCTTGATGTCTATCAGGCGC<br>GTTTTGACCATCTTCGTCTGATTATTGAGCAAAATAATTTATATGTGGCCG<br>GGTTCGTTAATACGGCAACAAATACTTTCTACCGTTTTTCAGATTTTACAC<br>ATATATCAGTGCCCGGTGTGACAACGGTTTCCATGACAACGGACAGCAGT<br>TATACCACTCTGCAACGTGTCGCAGCGCTGGAACGTTCCGGAATGCAAAT<br>CAGTCGTCACTCACTGGTTTCATCATATCTGGCGTTAATGGAGTTCAGTG<br>GTAATACAATGACCAGAGATGCATCCAGAGCAGTTCTGCGTTTTGTCACT<br>GTCACAGCAGAAGCCTTACGCTTCAGGCAGATACAGAGAGAATTTCGTCA<br>GGCACTGTCTGAAACTGCTCCTGTGTATACGATGACGCCGGGAGACGTG<br>GACCTCACTCTGAACTGGGGGCGAATCAGCAATGTGCTTCCGGAGTATC<br>GGGGAGAGGATGGTGTCAGAGTGGGGAGAATATCCTTTAATAATATATCA<br>GCGATACTGGGGACTGTGGCCGTTATACTGAATTGCCATCATCAGGGGG<br>CGCGTTCTGTTCGCGCCGTGAATGAAGAGAGTCAACCAGAATGTCAGATA<br>ACTGGCGACAGGCCCGTTATAAAAATAAACAATACATTATGGGAAAGTAAT<br>ACAGCTGCAGCGTTTCTGAACAGAAAGTCACAGTTTTTATATACAACGGG<br>TAAATAAAGGAGTTAAGCATGAAGAAGATGTTTATGGCGGTTTTATTTGCA<br>TTAGCTTCTGTTAATGCAATGGCGGCGGATTGTGCTAAAGGTAAAATTGA<br>GTTTTCCAAGTATAATGAGGATGACACATTTACAGTGAAGGTTGACGGGA<br>AAGAATACTGGACCAGTCGCTGGAATCTGCAACCGTTACTGCAAAGTGCT |

| | |
|---|---|
| | CAGTTGACAGGAATGACTGTCACAATCAAATCCAGTACCTGTGAATCAGG<br>CTCCGGATTTGCTGAAGTGCAGTTTAATAATGACTGA |
