## Supplementary Figures for "Protocell Arrays for Simultaneous Detection of Diverse Analytes"

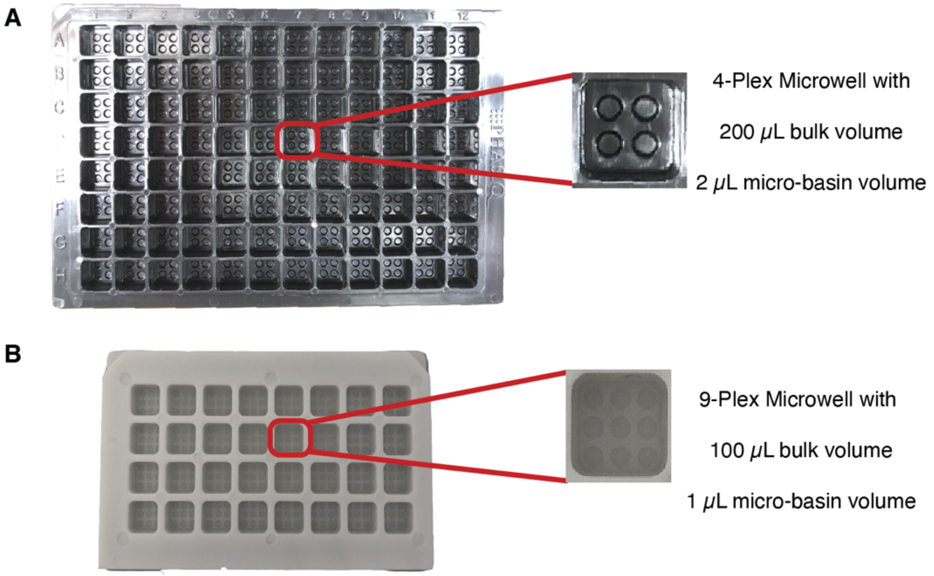


**Figure S1:** Custom-made polystyrene plates used for multiplexed protocell array reactions. Microwell layout and spacing are designed based on 96-well microplate standards from the Society for Laboratory Automation and Screening. **(A)** Black microwell plate with 2x2 arrays of 2 $\mu$L volume micro-basins for protocell placement. **(B)** White microwell plate with 3x3 arrays of 1 $\mu$L volume micro-basins for protocell placement.

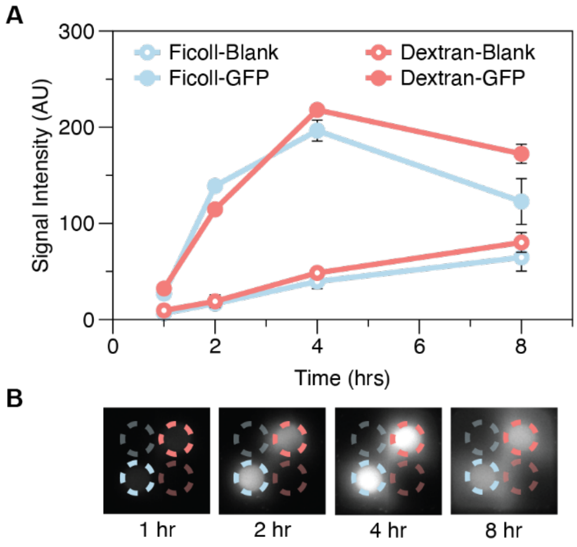


**Figure S2:** Time course characterization of CFE protein production and reaction compartmentalization in PEG-Ficoll and PEG-dextran ATPS. **(A)** Quantification of protocells containing CFE reactions with and without plasmid encoding for GFP expression. Both ATPS conditions exhibited similar fluorescence values at the detection time reported in the main text (3 hours), but the PEG-Ficoll system produces GFP slightly faster than the PEG-dextran system. Beyond 4 hours, phase separation of both PEG-Ficoll and PEG-dextran systems starts to deteriorate, resulting in greater fluorescent signal leakage to the surrounding environment, making the higher production by dextran-formed protocells less important. Disruption of the biphasic separation could be caused by polymer degradation in an increasingly acidic environment caused by cell-free protein systhesis[1]. Additionally, typical cell-free sensing reactions are on the order of 1 to 3 hours, making the results for longer time scales less relevant. Error bars represent standard deviations of technical triplicates. **(B)** Representative time-course fluorescence image of compartmentalized cell-free reactions in protocell arrays. Blue circles indicate Ficoll-formed protocell with a plasmid coding for GFP expression (bright blue) and without plasmid (faded blue). Red circles indicate dextran-formed protocell with a plasmid coding for GFP expression (bright red) and without plasmid (faded red).

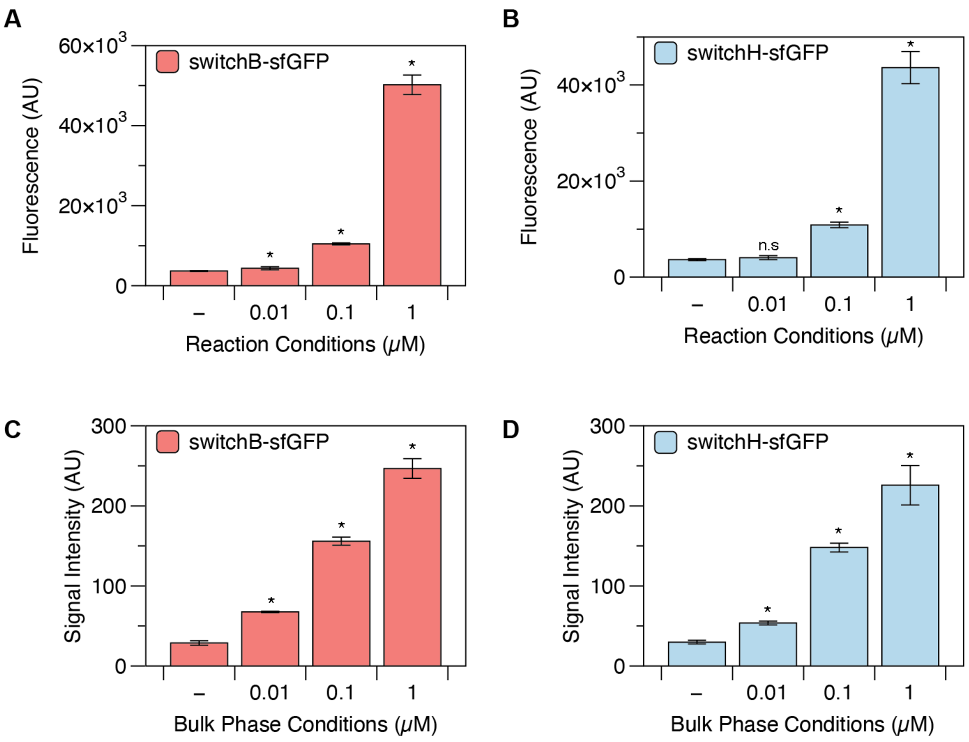


**Figure S3:** Comparison of toehold switch sensitivity to RNA triggers in CFE and protocell arrays. Reactions were incubated at 37 ˚C for 3 hours before fluorescent measurement on a plate reader (for CFE reactions) or a fluorescence imager (for protocell arrays). Compared to CFE sensors, protocell arrays show similar sensitivity and improved fold activation at lower RNA concentrations (0.01 μM). (**A-B**) Toehold switch B and H (respectively) activation at various RNA trigger concentrations in CFE reactions. (**C-D**) Toehold switch B and H (respectively) activation at various RNA trigger concentrations in protocell arrays. Statistical significance relative to basal expression is calculated using a two-tailed Student's t-test (n=3). Asterisks (*) indicate p-values < 0.05, n.s indicates no statistical significance. Error bars represent standard deviations of technical triplicates.

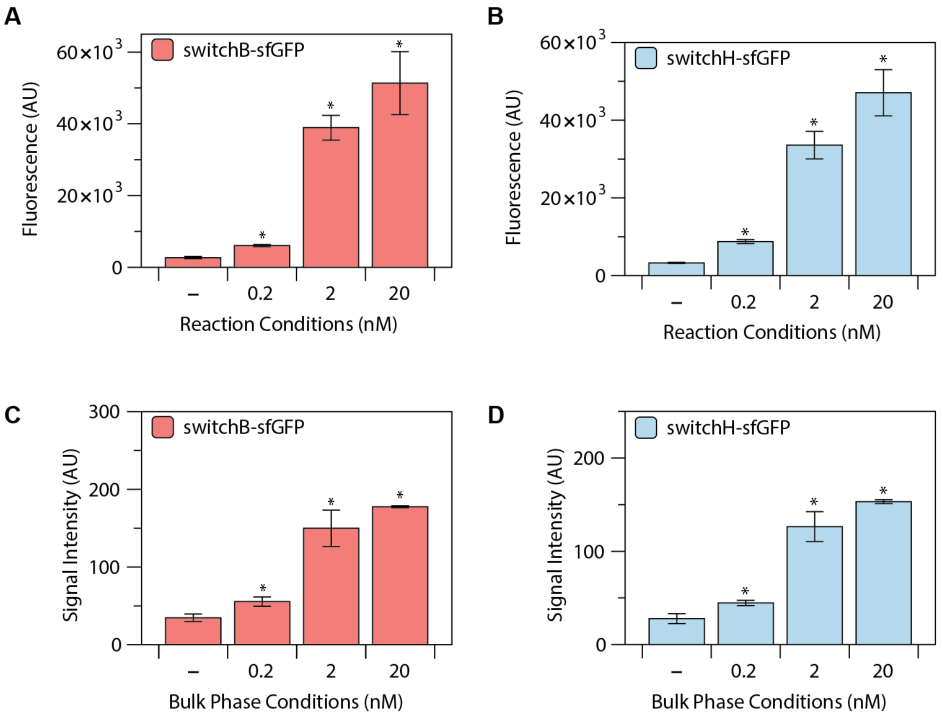


**Figure S4:** Comparison of toehold switch sensitivity to trigger-encoding linear DNA in CFE and protocell arrays. Reactions were incubated at 37 ˚C for 3 hours before fluorescent measurement on a plate reader (for CFE reactions) or a fluorescence imager (for protocell arrays). Compared to CFE sensors, protocell arrays show similar sensitivity. **(A-B)** Toehold switch B and H (respectively) activation at various trigger-encoding DNA concentrations in CFE reactions. **(C-D)** Toehold switch B and H (respectively) activation at various trigger-encoding DNA concentrations in protocell arrays. Statistical significance relative to basal expression is calculated using a two-tailed Student's t-test (n=3). Asterisks (*) indicate p-values < 0.05. Error bars represent standard deviations of technical triplicates.

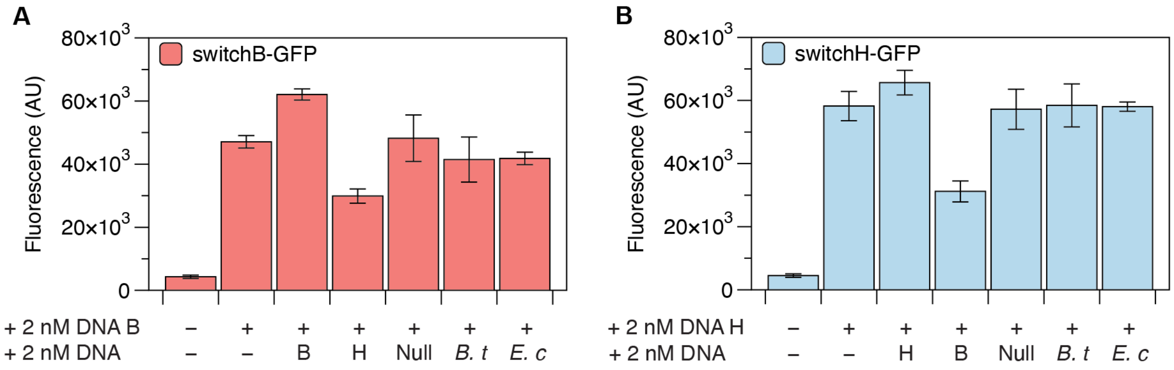


**Figure S5:** Co-expression of triggers B and H in CFE reactions mutually represses their output. This inhibition effect is specific to the combination of triggers B and H, as evidenced by the lack of repression when random (Null) and previously characterized trigger sequences (*Bacteroides thetaiotaomicron* and *Escherichia coli, B. t* and *E. c* respectively)[3] are also added to the CFE reactions. This effect occurs for any CFE sensing reaction, not just in protocell arrays. (**A**) Trigger B was co-expressed with different linear DNA transcribing RNA triggers. Only the addition of trigger H significantly reduced GFP production from the switchB-GFP plasmid. **(B)** Trigger H was co-expressed with different linear DNA transcribing RNA triggers, and only the addition of trigger B reduced GFP production from the switchH-GFP plasmid.

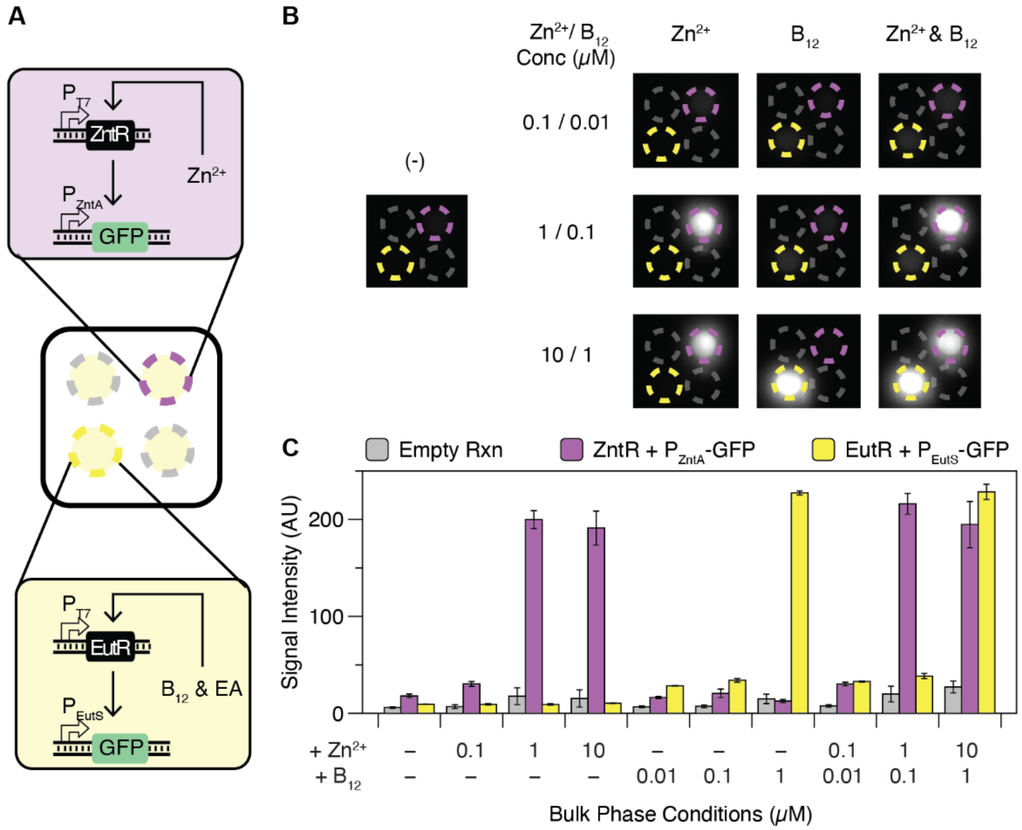


**Figure S6: (A)** Schematic of protocell array setup for multiplexed analysis of zinc and vitamin $B_{12}$. Zinc modulates GFP expression by binding to the transcriptional regulator ZntR, which in turn activates transcription from its cognate promoter $P_{ZntA}$ (purple schematic). Vitamin $B_{12}$ modulates GFP expression by binding to the transcriptional regulator EutR with ethanolamine (EA) as a co-factor. EutR then activates transcription from its cognate promoter $P_{EutS}$ (yellow schematic). Gray circles represent negative control protocells with no plasmid DNA. **(B)** Representative fluorescence images of small molecule multiplexing results at different input and concentration conditions. (-) indicates neither zinc nor vitamin $B_{12}$ was added in the bulk phase. Zinc was tested at 0.1, 1, and 10 $\mu$M. Vitamin $B_{12}$ was tested at 0.01, 0.1, and 1 $\mu$M. All micro-wells containing vitamin $B_{12}$ in the bulk phase also had 50 $\mu$M EA as a cofactor for sensor activation. **(C)** Quantification of fluorescence images in (B) and their replicates. Error bars represent standard deviations of technical triplicates.

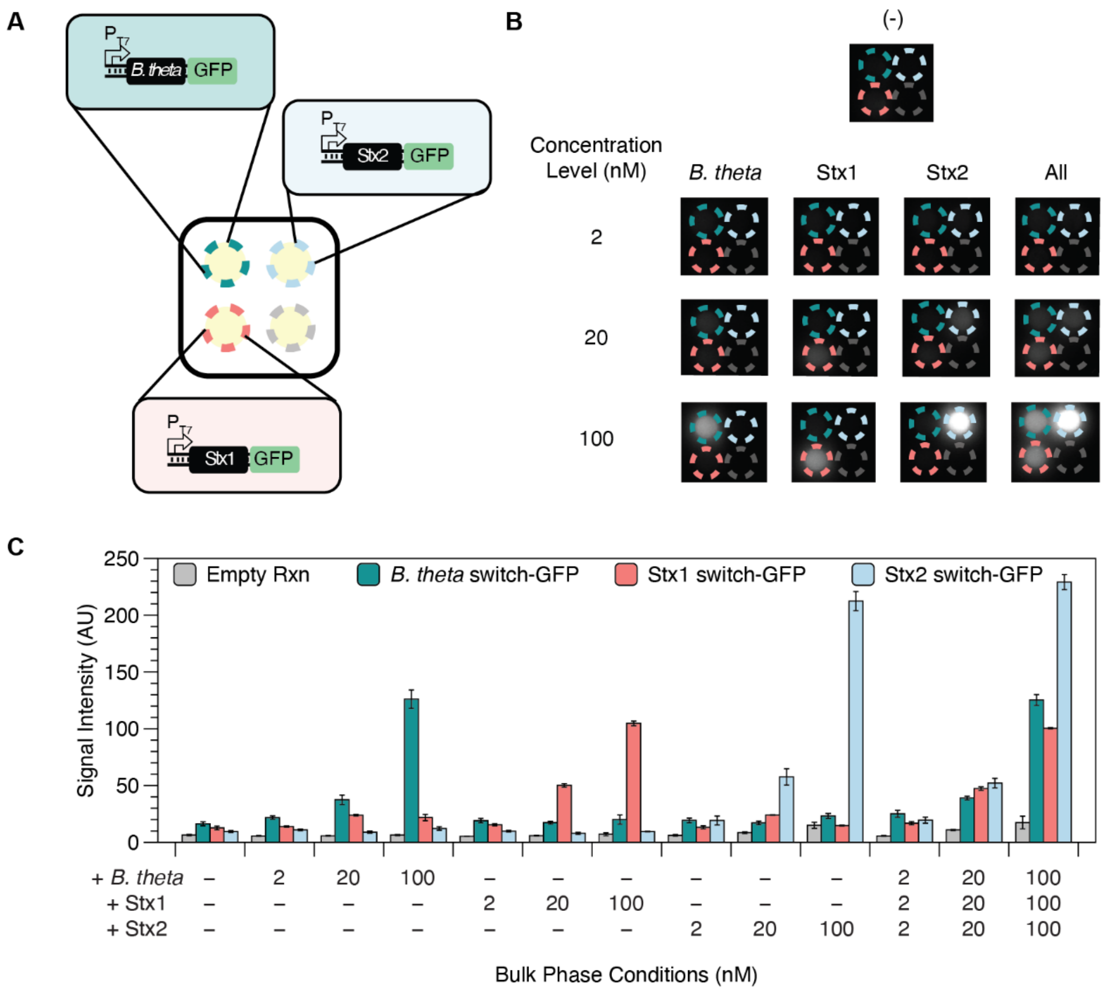


**Figure S7: (A)** Schematic of protocell array setup for multiplexed detection of *B. theta* and STEC bacteria. Teal, red, and blue circles represent micro-basins containing *B. theta*, Stx1, and Stx2 protocell sensors, respectively. The gray circle represents a negative control protocell with no plasmid DNA. **(B)** Representative fluorescence image of multiplexed nucleic acid sequence detection results at different input and concentration conditions. Linear DNA for expression of triggers was amplified from target-specific primers and added to the bulk phase at concentrations of 2, 20, and 100 nM. **(C)** Quantification of fluorescence images from (B) and their replicates. Error bars represent standard deviations of technical triplicates.

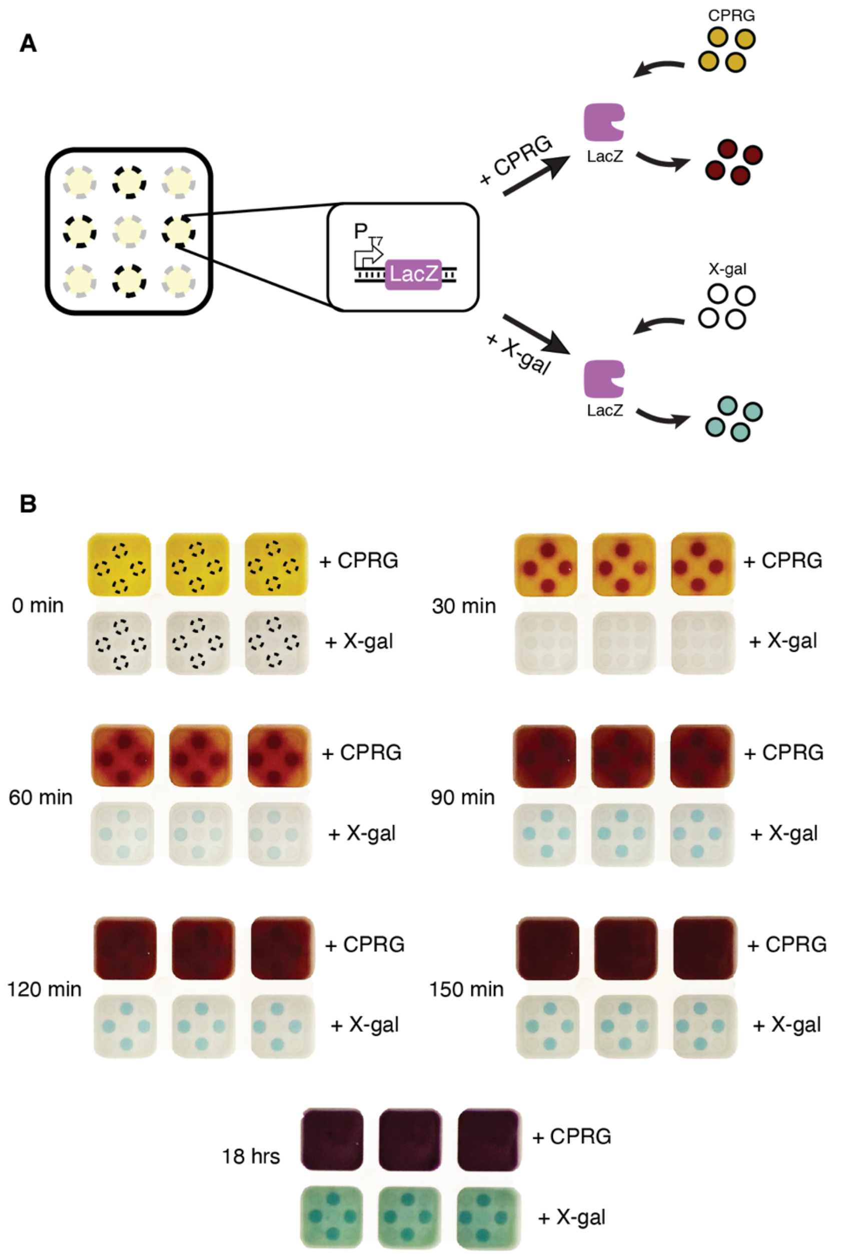


**Figure S8**: Protocell array output can be interpreted without equipment. **(A)** Schematic of visually interpretable protocell array sensing reactions. Black circles represent micro-basins containing protocells that constitutively produce LacZ. The bulk phase contains either CPRG (yellow in panel (B)) or X-gal (colorless in panel (B)) as the substrate for pigment production. Once produced, the LacZ enzyme either cleaves CPRG to form chlorophenol red (CPR, red) or cleaves X-gal to form a blue precipitate. **(B)** Time course pigment production from two substrates. Bulk phase containing CPRG yields visible color change within 30 minutes of incubation, but the pigment readily diffuses into the bulk phase at later time points, making test results uninterpretable. Bulk phase containing X-gal produces visible color more slowly, but the color remains localized over the entire incubation period, and still mostly localized even after overnight incubation.

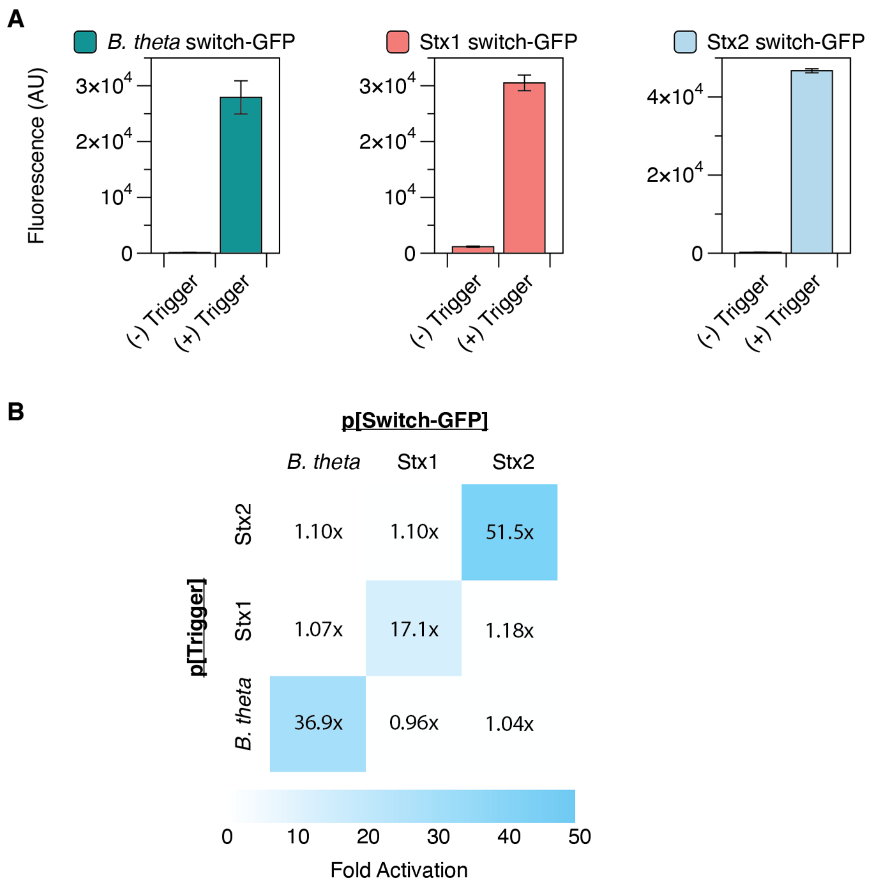


**Figure S9: (A)** Initial validation of *B. theta*, Stx1, and Stx2 switch activation in response to their cognate triggers in CFE reactions. All triggers and switches were expressed from plasmids. Error bars represent standard deviations of technical triplicates. **(B)** Specificity assessment of previously developed *B. theta* toehold switch and newly developed Stx1 and Stx2 toehold switches. Only correctly paired triggers and switches showed high levels of GFP activation.

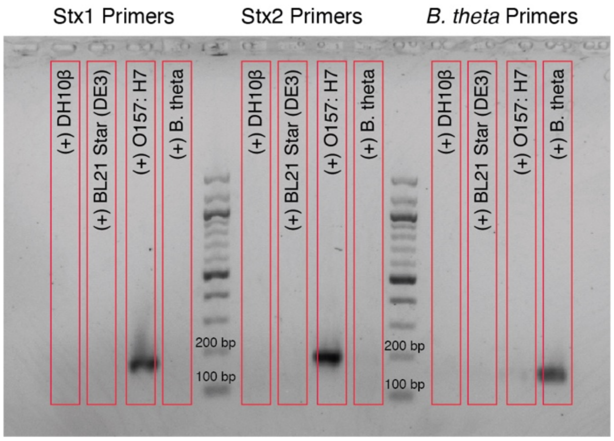


**Figure S10:** Validation of target-specific trigger amplification. Primers were designed to amplify either Stx1, Stx2 or *B. theta* triggers. Target DNA was amplified when the appropriate template was added (STEC O157:H7 genomic DNA for Stx1 and Stx2, and *B. thetaiotaomicron* genomic DNA for *B. theta*). No amplification was observed on the genomic DNA of common lab *E. coli* strains DH10β or BL21 Star (DE3). PCR products were run on a 2% agarose gel.

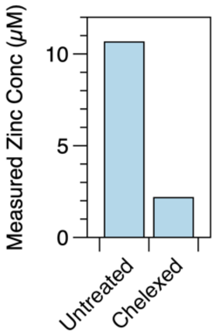


**Figure S11.** ICP-MS measurement of zinc concentrations in untreated and Chelex-100 treated human serum.
